# Structural and Functional Insights into the Evolution of SARS-CoV-2 KP.3.1.1 Spike Protein

**DOI:** 10.1101/2024.12.10.627775

**Authors:** Ziqi Feng, Jiachen Huang, Sabyasachi Baboo, Jolene K. Diedrich, Sandhya Bangaru, James C. Paulson, John R. Yates, Meng Yuan, Ian A. Wilson, Andrew B. Ward

## Abstract

The JN.1-sublineage KP.3.1.1 recently emerged as the globally prevalent SARS-CoV-2 variant, demonstrating increased infectivity and antibody escape. We investigated how mutations and a deletion in the KP.3.1.1 spike protein (S) affect ACE2 binding and antibody escape. Mass spectrometry revealed a new glycan site at residue N30 and altered glycoforms at neighboring N61. Cryo-EM structures showed that the N30 glycan and rearrangement of adjacent residues did not significantly change the overall spike structure, up-down ratio of the receptor-binding domains (RBDs), or ACE2 binding. Furthermore, a KP.3.1.1 S structure with hACE2 further confirmed an epistatic effect between F456L and Q493E on ACE2 binding. Our analysis shows SARS-CoV-2 variants that emerged after late 2023 are now incorporating reversions to residues found in other sarbecoviruses, including the N30 glycan, Q493E, and others. Overall, these results inform on the structural and functional consequences of the KP.3.1.1 mutations, the current SARS-CoV-2 evolutionary trajectory, and immune evasion.

## Introduction

After five years of evolution in the human population, SARS-CoV-2 continues to circulate globally, albeit with reduced severity and mortality. The continuous evolution has led to the emergence of a plethora of variants with multiple mutations, especially on the spike protein, allowing them to evade antibodies induced by prior infection or vaccination and/or enhance viral fitness and transmissibility. By the start of 2024, the JN.1 variant had outcompeted the previous XBB lineages^1–4^. Subsequently, the JN.1-derived subvariants (e.g., KP.2, KP.3) became the dominant strains^5–7^. Recently, KP.3.1.1 emerged as the predominant SARS-CoV-2 variant circulating worldwide, overtaking its parent lineage KP.3 and earlier KP.2 variants^8–12^. The proportion of KP.3.1.1 in the population continued to increase, and by September 2024, accounted for more than 50% of global cases^13,14^.

KP.3.1.1 has acquired a higher relative effective reproduction number (R_e_), higher infectivity, and greater neutralization escape than KP.3 and previous variants^8–12,15^. Compared to JN.1, both KP.3 and KP.3.1.1 share two mutations F456L and Q493E in the receptor-binding domain (RBD) of the spike protein. These mutations work together to facilitate further antibody escape and exhibit better binding to the host receptor, human angiotensin-converting enzyme 2 (hACE2)^16,17^. An S31 deletion in the N-terminal domain (NTD) of KP.3.1.1 at ^30^NSFT^33^ introduces a potential N-linked glycosylation site (PNGS, NxS|T motif) at N30 (NFT) that was previously observed in some non-SARS-CoV-2 sarbecoviruses. Previous studies have suggested that the S31 deletion and potential N30 glycosylation may alter the local spike conformation and enable the RBD to adopt a more “down” conformation, potentially enabling escape from certain classes of pre-existing neutralizing antibodies^9,11^. A V1104L mutation, located on the lower stem of the spike S2, in KP.3.1.1 as well as in earlier subvariants (such as KP.2, KP.2.3, and KP.3) was considered to have a minimal effect on antibody evasion^18^. Currently, little is known about the structural effects of the deletion and N30 glycosylation and the consequences and mechanism of immune escape in KP.3.1.1.

In this study, we used single-particle cryo-electron microscopy (cryo-EM) to investigate structures of the apo KP.3.1.1 spike and its complex with hACE2. By analyzing the molecular interactions between the KP.3.1.1 RBD and hACE2, we rationalized the epistatic and synergistic effects of F456L and Q493E mutations, which markedly enhanced the binding affinity to hACE2 over the Q493E mutation alone, similar to the F456L/Q493E synergy previously revealed in the KP.3 variant^17^. In addition, to dissect how the mutations S31Δ, F456L, and Q493E affect the overall spike stability, we introduced these mutations into the JN.1.11 spike protein. The ratio of RBD “up” and “down” in these variants was analyzed semi-quantitatively through cryo-EM. Through biochemical and structural analysis by mass spectrometry and cryo-EM, we demonstrated that the S31 deletion enabled N-glycosylation at N30 and caused a side-chain 180° flip of the adjacent F32 residue. The S31 deletion and N30 glycosylation exhibited minimal effect on the spike protein conformation, and did not affect spike binding to hACE2 or to the broadly neutralizing antibody (bnAb) BD55-1205^19^. However, addition of N-glycans at N30 altered the glycoforms at a spatially adjacent N-glycosylation site at residue N61. Through bioinformatic analysis, we also observed an important trend in recent SARS-CoV-2 evolution—many mutations in recent variants of concern recapitulate residues found in other sarbecoviruses, including but not limited to Q493E and the PNGS at N30. Most of these re-occurring mutations have emerged since late 2023. Overall, these findings advance our understanding of current SARS-CoV-2 evolution and provide new insights for prediction of subsequent evolution, as well as the design of vaccines and therapeutic antibodies against current and future SARS-CoV-2 variants.

## Results

### Deletion of conserved S31 on the NTD introduces N30 glycosylation

Compared to the JN.1 spike protein, KP.3 carries two mutations in the RBD (F456L and Q493E), and one mutation in the S2 domain (V1104L). KP.3.1.1 contains only one additional change compared to KP.3, a deletion in the NTD (S31Δ) (**Figure 1A**). JN.1.11 is JN.1 plus V1104L and JN.1.11.1 is JN.1.11 plus F456L. Sequence alignment of sarbecoviruses showed that S31 is conserved in all sarbecoviruses (**Figure 1B**). The S31Δ changes the NTD sequence from ^30^NSFT^33^ to NFT, and introduces a potential N-glycosylation site at N30, which has been previously observed by cryo-EM in certain bat and pangolin sarbecovirus strains^20–22^. However, in KP.3.1.1, it is a rare instance of an “NxT” sequon at N30, compared to a few other sarbecoviruses with an “NxS” sequon (**Figure 1B**). According to the data from GISAID, KP.3.1.1 has outcompeted other JN.1-derived variants (e.g., JN.1.11.1, KP.2, KP.2.3, KP.3, LB.1, etc.) and become the dominant strain in the United States and globally as of November, 2024 ^13,14^ (**Figure 1C**). Thus, the deletion at S31 and the ensuing glycosylation site at N30 appear to endow a fitness advantage over its most recent predecessors.

**Figure 1.**
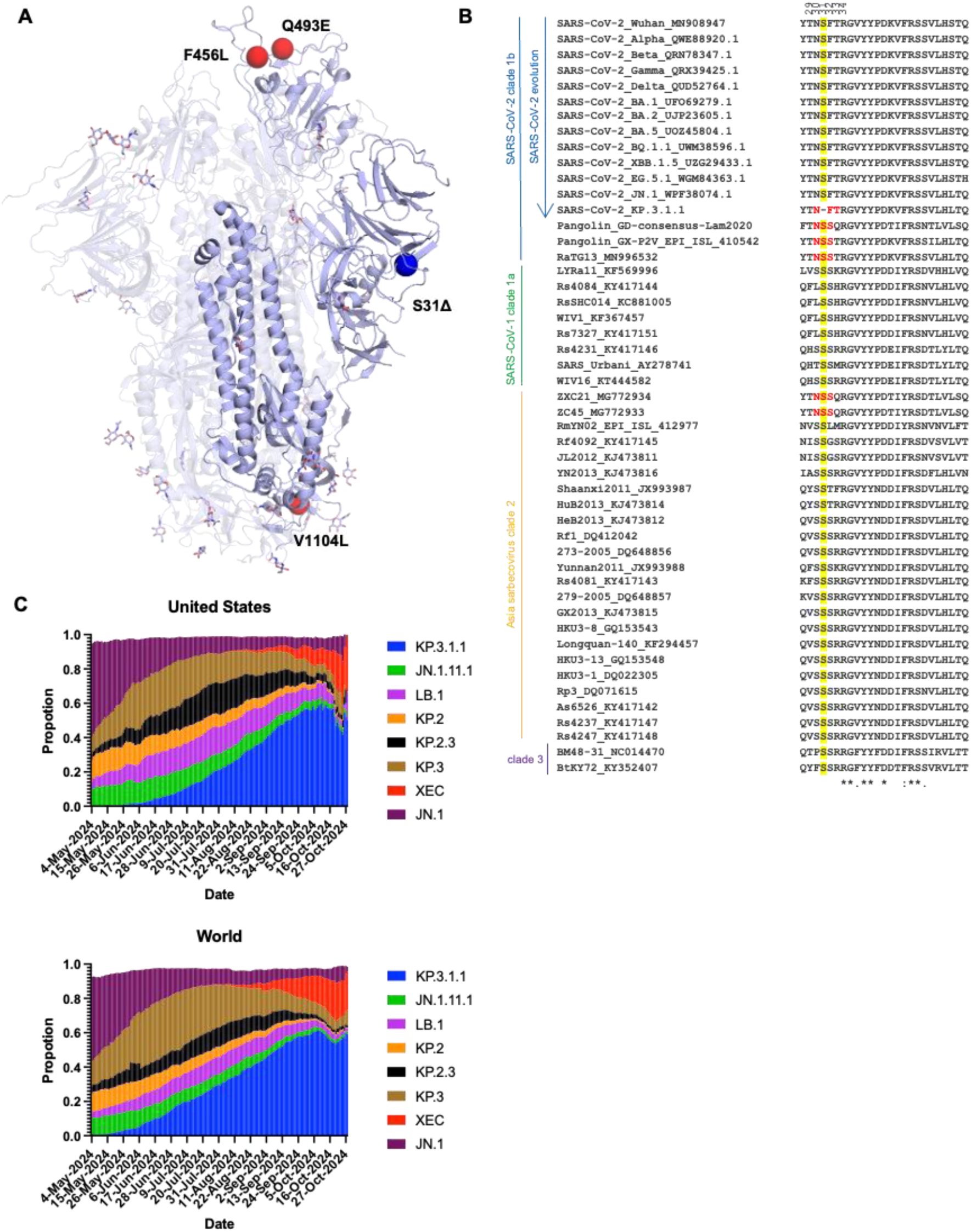
Deletion of conserved S31 on the NTD introduces an N30 glycosylation site. **(A)** Mutations (red) and a deletion (blue) in KP.3.1.1 in the context of JN.1 S (lavender) (PDB ID: 8Y5J)^1^. **(B)** Sequence alignment of sarbecoviruses including several SARS-CoV-2 Variants of Concern (VoC). Yellow highlights the conserved S31 across sarbecoviruses except for KP.3.1.1. Red shows the N-glycosylation sequon at N30. Residues identical in all aligned sequences are labeled by an asterisk (*), whereas a colon (:) and a period (.) indicate strongly similar and less similar sequences, respectively. The sequence alignment was performed with Clustal Omega^48^. **(C)** Frequencies of circulating variants JN.1, JN.1.11.1, KP.2, KP.2.3, KP.3, KP.3.1.1, and LB.1 sequences in the United States and worldwide from 05/04/2024 to 11/01/2024. Data were collected from covSPECTRUM^13^.

To validate the potential N-glycosylation site at N30, we utilized the DeGlyPHER^23^ approach, which combines sequential deglycosylation with bottom-up mass spectrometry-based proteomics to identify N-glycosylation sites and characterize site-specific glycan heterogeneity across seven spike protein variants including JN.1, JN.1.11, JN.1.11.1, and KP.3.1.1, as well as engineered variants JN.1.11+S31Δ, JN.1.11.1+S31Δ, JN.1.11+Q493E+S31Δ (**Figure 2A**). Our results demonstrate that S31Δ indeed induces a completely glycan-occupied N30 glycosylation site in all of the variants carrying the S31 deletion, including JN.1.11+S31Δ, JN.1.11.1+S31Δ, JN.1.11+Q493E+S31Δ, and KP.3.1.1. The N30 site (NxT motif) was found to be predominantly occupied with high mannose glycans. These findings confirm the previous hypothesis that the S31 deletion would introduce a PNGS at N30^9–11^.

**Figure 2.**
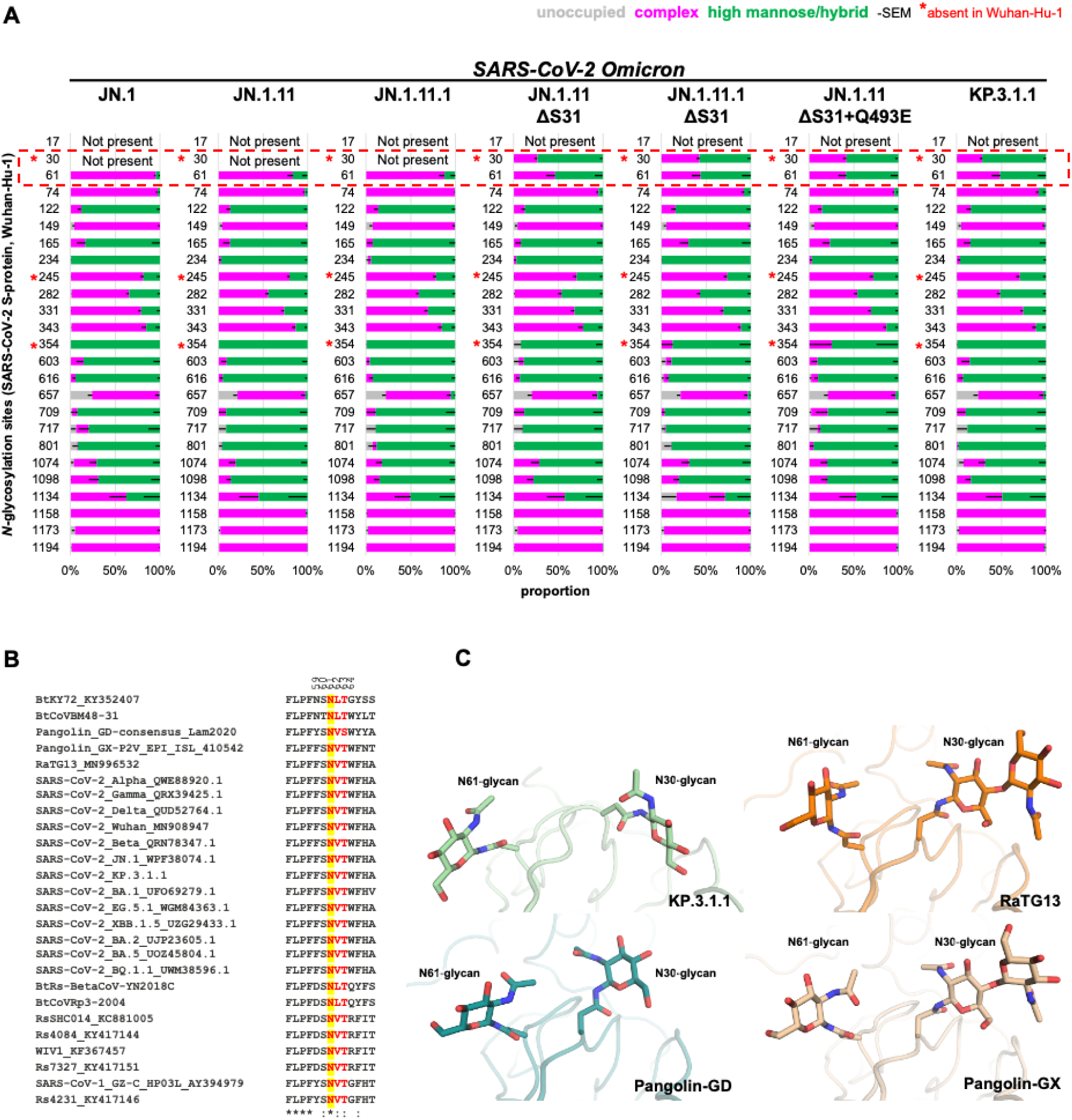
Analysis of the N-glycan landscape on the spike proteins. **(A)** Bar-graphs demonstrating site-specific differences in glycosylation, if any, between JN.1 and six natural or engineered mutants (JN.1.11, JN.1.11.1, JN.1.11+S31Δ, JN.1.11.1+S31Δ, JN.1.11+S31Δ+Q493E, and KP.3.1.1) as determined by DeGlyPHER. *N*-glycosylation states (unoccupied, complex, or high mannose) are color-coded. Error bars represent mean–SEM. S31Δ resulted in glycosylation at N30 and a concomitant significant change of complex to high mannose *N*-glycans at N61 (*p* = 0.002; KP.3.1.1 vs. JN.1). **(B)** Sequence alignment of sarbecoviruses including several Variants of Concern (VoC) of SARS-CoV-2. Yellow highlights the conservation of N61 across all sarbecoviruses and red shows the N-glycosylation sequon leading to glycosylation at N61. Residues identical in all aligned sequences are labeled by an asterisk (*), whereas a colon (:) and a period (.) indicate strongly similar and less similar sequences, respectively. The sequence alignment was performed with Clustal Omega^48^. **(C)** Zoomed-in view of the close proximity and interaction between the N30 and N61 glycans in KP.3.1.1, RaTG13 (PDB ID: 6ZGF)^21^, Pangolin-CoV-GD (PDB ID: 7BBH)^20^, and Pangolin-CoV-GX (PDB ID: 7CN8)^22^.

### Structural overview of the Omicron KP.3.1.1 other JN.1-derived mutants spike proteins

To investigate the effects of N30 glycosylation introduced by S31Δ and two RBD mutations (F456L and Q493E) on the structure and function of the spike protein, we determined the cryo-EM structures of the recent SARS-CoV-2 variants KP.3.1.1 and JN.1.11 along with engineered variants JN.1.11+S31Δ+Q493E, JN.1.11.1+S31Δ, and JN.1.11+S31Δ (**Figure 3, Table S1, Figure S1, and Figure S2**). All the spike proteins exhibited three conformational states: “One RBD Up”, “All RBD Down”, and “Flexible” (i.e., one or two RBDs disordered in the density maps due to flexibility). Similar conformational states (“one up” and “all down”) were observed in the spike protein structures of KP.3.1.1’s ancestors, including BA.2, BA.2.12.1, BA.2.75, and BA.2.86, which showed both “One RBD Up” and “All RBD Down” conformations^24–26^. Prior studies also indicated that the JN.1 spike protein exhibited either a “One RBD Up” or “All RBD Down” conformation^1,27^. Using 3D variability analysis, we quantified the particle proportion for each conformational state and found that S31Δ did not affect the proportions of “One RBD Up” and “All RBD Down” particles by comparing the proportion of JN.1.11 and other spike proteins that have S31Δ (**Figure 3G and Figure S1**). Additionally, S31Δ did not markedly affect the thermostability of the spike protein compared to JN1.11, JN1.11.1, and engineered variants as assessed by nanoDSF, but retained the substantially increased Tm over WT SARS-CoV-2 (Tm of 46 vs 32 **°**C) (**Figure S3**). Structural alignment of SARS-CoV-2 WT, JN.1, and JN.1.11 spike proteins with KP.3.1.1 showed that NTD and RBD exhibited greater RMSDs than S2 (**Figure S4**). V1104L has little effect on conformational changes; V1104 and L1104 are both surrounded by a conserved hydrophobic pocket formed by P1090, F1095, I1115, and T1120 in the same protomer (**Figure S5**).

**Figure 3.**
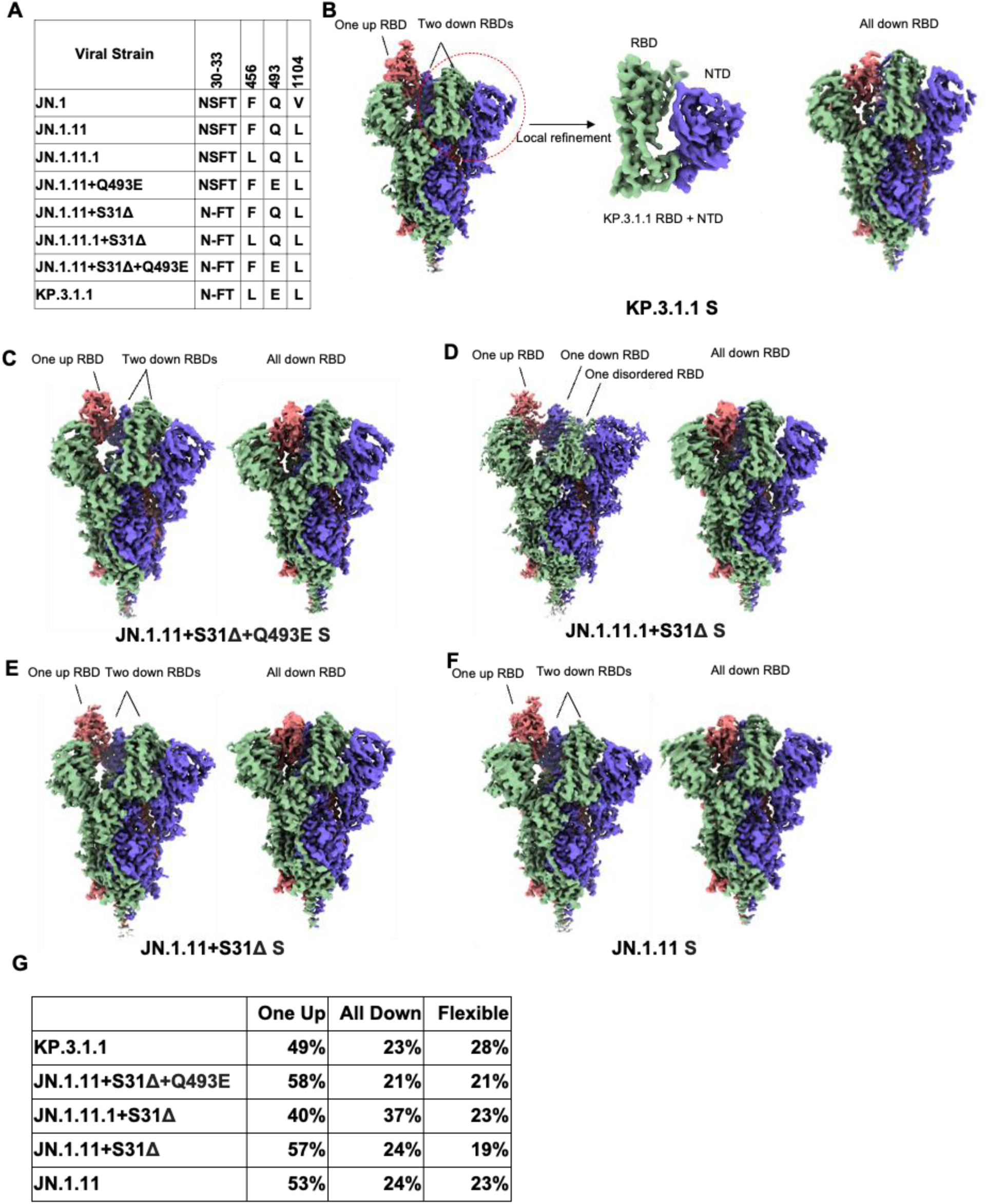
Structural overview of Omicron KP.3.1.1 and other JN.1-derived mutants spike proteins. **(A)** Sequence alignment showing specific mutated amino acids in KP.3.1.1 and other naturally occurring or engineered JN.1-derived spike mutants. **(B)** Cryo-EM density maps of KP.3.1.1 S (“One RBD Up” on the left and “All RBD Down” on the right) and map around KP.3.1.1 RBD+NTD (middle) after local refinement. Maps are colored by chain with the three protomers of the spike trimer in blue, green, and salmon. **(C-F)** Side view of the cryo-EM density maps of JN.1.11+S31Δ+Q493E, JN.1.11.1+S31Δ, JN.1.11+S31Δ, JN.1.11 S trimers, respectively. Maps are colored by chain as in B. Left, S in the “One RBD Up” conformation. Right, S in the “All RBD Down” conformation. **(G)** The percentage of three conformational states (one RBD up, all RBD down, and flexible (i.e., one or two RBDs disordered in the density maps due to flexibility), observed for the spike proteins in **B-F**. After 3D variability analysis, particles are clustered into different groups by their conformational state. We then counted the state of each cluster and the number of corresponding particles. (See also figures S1, S2, S3, S4, S5, and Table S1).

### Molecular basis of KP.3.1.1 S binding to hACE2

To examine whether S31Δ and the other recent mutations affect S-hACE2 binding, we used bio-layer interferometry (BLI) to test the binding affinities of various spike proteins to hACE2 (**Figure 4F and Figure S6**). The data from three paired variant sets (JN.1.11 & JN.1.11+S31Δ, JN.1.11.1 & JN.1.11.1+S31Δ, and JN.1.11+Q493E & JN.1.11+Q493E+S31Δ) showed that S31Δ has little impact on hACE2 binding, which is consistent with previous studies^10,28^. This result is also concordant with our observation that S31Δ and the resulting N30 glycosylation do not interfere with the “Up/Down” configuration of spike RBD, where the “up” conformation of RBD is required for binding to hACE2. Besides, while F456L alone did not affect hACE2 binding (JN.1.11.1 has F456L compared to JN.1.11), Q493E alone substantially reduced binding affinity (**Figure 4F**), which is also consistent with prior research and is likely due to the electrostatic repulsion between hACE2-E35 and RBD-E493^17^ (**Figure 4D**).

**Figure 4.**
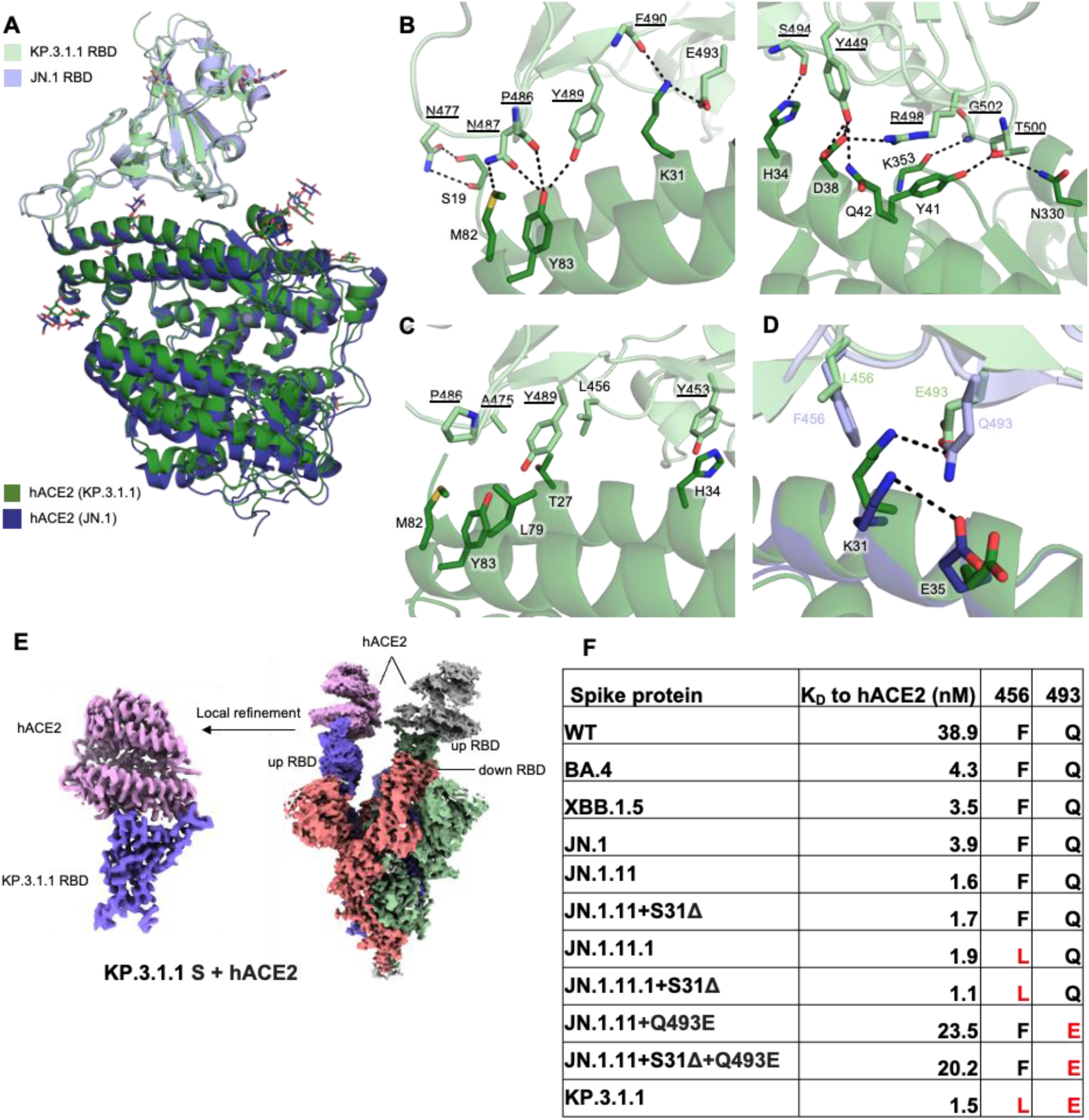
Molecular interactions of KP.3.1.1 RBD with hACE2. **(A)** Structural alignment of KP.3.1.1 RBD/hACE2 with JN.1 RBD/hACE2 (PDB ID: 8Y18)^1^ from cryo-EM structures of their spike proteins with hACE2 using RBD for alignment. Cα RMSD between the two RBDs is 0.9 Å. KP.3.1.1 RBD and hACE2 are colored in light green and green, respectively. JN.1 RBD and its hACE2 are colored in light blue and blue, respectively. **(B)** Molecular details of hydrophilic interactions between KP.3.1.1 RBD and hACE2. Conserved residues between KP.3.1.1 and JN.1 are underlined. Residues are numbered according to their positions on the SARS-CoV-2 spike protein sequence. Hydrogen bonds and salt bridges are represented by black dashed lines. **(C)** Hydrophobic interactions between KP.3.1.1 RBD and hACE2. Aromatic and aliphatic residues are displayed. **(D)** Structural analysis of cooperative interactions between F456L, Q493E, and ACE2. F456 and Q493 in JN.1 RBD, and L456 and E493 in KP.3.1.1 are displayed. Two salt bridges between K31 and E35 in the JN.1 RBD/hACE2 complex (intrachain), and between K31 and E493 (interchain) in the KP.3.1.1 RBD/hACE2 complex, are shown as black dashed lines. **(E)** Left, cryo-EM density map corresponding to the KP.3.1.1 RBD/hACE2 interaction from focused local refinement of cryo-EM structure with spike protein and hACE2. Right, cryo-EM density of the entire KP.3.1.1 S/hACE2 complex. Maps are colored by chain with the three protomers of the spike trimer in blue, green, and salmon and hACE2 in pink and grey. **(F)** Binding affinity of hACE2 to recombinant spike proteins. Binding affinity is expressed as the nanomolar dissociation constant (*K*_D_). (See also figure S6).

To investigate the interactions between KP.3.1.1 S and the cell receptor, we also determined a structure of KP.3.1.1 S/hACE2 complex at 2.9 Å resolution using cryo-EM. Two RBDs were found to be in the “up” position with each RBD binding to an hACE2 molecule, while the third RBD is in the “down” position (**Figure 4E**). The local refinement map of KP.3.1.1 RBD with hACE2 was also excellent and resolved at 3.1 Å resolution (**Figure 4E, Table S1, Figure S2B**). Structural alignment with the JN.1 RBD-hACE2 complex (PDB: 8Y18) revealed similar binding (Cα RMSD = 0.9 Å)^1^ (**Figure 4A**). Residues on the receptor-binding site (RBS) of KP.3.1.1 RBD are highly conserved with JN.1 and form extensive electrostatic interactions with hACE2: E493 and R498 form salt bridges with K31 and D38, respectively; the main-chain of P486, F490, S494, and G502 form H-bonds with the side-chains of Y83, K31, H34, and K353, respectively; N477, and Y489 H-bond with S19, and Y83, respectively; Y449 H-bonds with D38 and N42, N487 with M82 and Y83, and T500 with Y41 and N330 (**Figure 4B**). Several conserved aromatic and aliphatic residues (Y453, A475, P486, and Y489) also establish hydrophobic interactions with hACE2 (**Figure 4C**). Q493E alone reduced ACE2 binding, while F456L alone had a minimal impact on binding; however, ACE2 binding with the Q493E mutation was recovered in the context of F456L (**Figure 4F and Figure S6, JN.1.11+S31Δ+Q493E vs KP.3.1.1**). This phenomenon has been observed and structurally depicted in the KP.3 variant^17^. Thus, we also observed the epistatic effect of F456L and Q493E in KP.3.1.1, which together enhanced hACE2 binding, compared to the substantially reduced binding with Q493E alone^16,17^. RBD-Q493E alone in the presence of F456 would likely be disfavored due to an electrostatic repulsion with ACE2-E35 (**Figure 4D, 4F, and Figure S6**). The bulkier RBD-F456 side chain shifts ACE2-K31 to form an intrachain salt-bridge interaction with ACE2-E35, thereby reducing its interaction with RBD E493. However, when the smaller side chain of L456 is present, K31 can move and form a salt bridge with E493 (**Figure 4D**)^10^ that is a stronger interaction than the hydrogen bond between K31 and Q493 in the presence of F456, which explains how F456L and Q493E can synergize to recover the binding affinity with hACE2.

### BD55-1205 binding to KP.3.1.1 and other S31Δ mutants

BD55-1205 is a “Class 1” antibody encoded by IGHV3-55/3-66 germline, showing potent and broad neutralizing ability across all variants from WT to JN.1^19^. The epitope residues of BD55-1205 mostly overlap with hACE2, explaining its potent neutralizing ability^19,29^. To determine whether KP.3.1.1 or other S31Δ variants escape BD55-1205, we used BLI to measure binding affinities between spike proteins and the Fab (**Figure 5G and Figure S7**). BD55-1205 retained strong binding to KP.3.1.1 S and other mutants, indicating that S31Δ had minimal effect on BD55-1205 binding, further supporting our previous finding that S31Δ did not alter the RBD “up/down” ratio. In addition, other spike mutants with Q493E retain BD55-1205 binding, while F456L resulted in a threefold decrease in binding affinity where the only mutation of JN.1.11.1 is F456L compared to JN.1.11 **(Figure 5G and Figure S7)**. We superimposed the structure of BD55-1205/XBB.1.5 RBD complex (PDB ID: 8XE9) to KP.3.1.1 S and the RBDs showed a highly similar conformation (Cα RMSD = 0.9 Å) (**Figure 5D**)^19^. The affinity difference between BD55-1205 and different JN.1 subvariants is similar to previous results^19^, and consistent with Q493E possibly forming a salt bridge with V_H_ R102 (**Figure 5E**), while F456L may reduce the hydrophobic interactions with V_H_ Y33 and V_L_W94 (**Figure 5F**). BD55-1205 forms extensive hydrophilic (H-bonds and salt bridges) interactions with the RBD. Furthermore, the epitope residues involved in hydrophilic interactions are highly conserved between XBB.1.5 and KP.3.1.1, explaining the nM binding affinity between BD55-1205 and the KP.3.1.1 spike protein (**Figure 5E**). However, hydrophobic interactions may be weakened by two mutations L455S and F456L (**Figure 5F**). The binding of BD55-1205 is substantially reduced to all of these variants compared to SARS-CoV-2 WT (**Figure 5G**).

**Figure 5.**
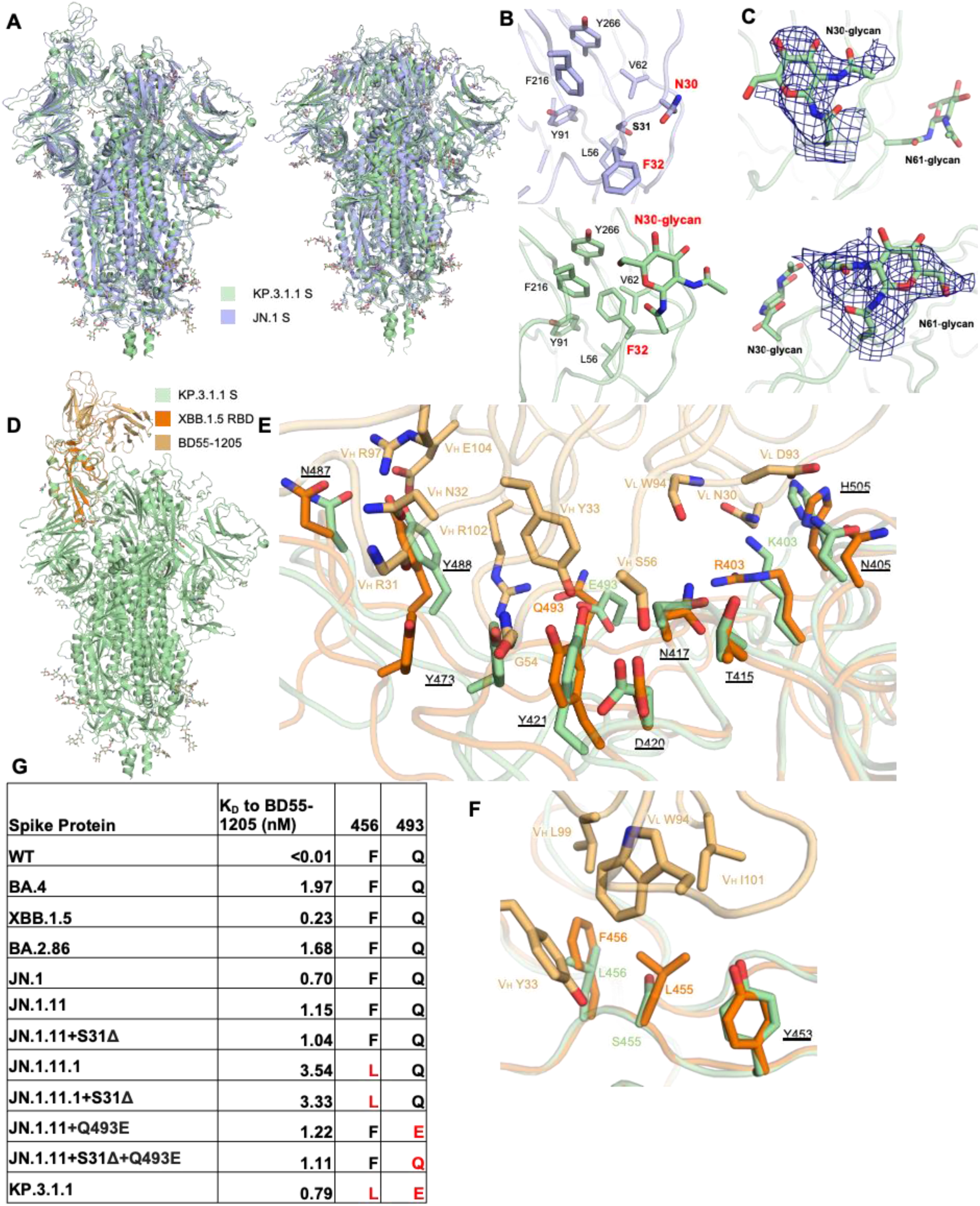
Structural details of S31Δ, N30 glycosylation, and BD55-1205 binding to KP.3.1.1 and other S31Δ mutants. **(A)** Superimposed cryo-EM structures of KP.3.1.1 S and JN.1 S in both “One RBD Up” (PDB ID: 8Y5J)^1^ and “All RBD Down” (PDB ID: 8X4H)^27^ states. Cα RMSD’s are 1.1 Å and 0.9 Å respectively. KP.3.1.1 S and JN.1 S are colored light green and light blue, respectively. **(B)** Zoomed-in view around the N30 glycan site. Compared to the JN.1 S, S31Δ results in an F32 sidechain flip of 180° to now fit into the hydrophobic pocket formed by L56, V62, Y91, F216, and Y266. N30 is glycosylated in KP.3.1.1 S. **(C)** Zoomed-in view of N30 and N61 glycans with the cryo-EM density map in deep blue. **(D)** Superimposed structures of KP.3.1.1 S with the XBB.1.5 RBD/BD55-1205 complex (PDB ID: 8XE9)^19^ to identify possible interactions of BD55-1205 with KP.3.1.1. KP.3.1.1 S, XBB.1.5 RBD, and BD55-1205 are colored in light green, orange, and light orange, respectively. **(E, F)** Zoomed-in view of the hydrophilic and hydrophobic interactions between XBB.1.5 RBD and BD55-1205, respectively, Corresponding residues in KP.3.1.1 in the binding site are also displayed. Conserved residues between KP.3.1.1 and XBB.1.5 are underlined. **(G)** Binding affinity of BD55-1205 Fab to recombinant spike proteins. Binding affinity is expressed as the nanomolar dissociation constant (*K*_D_). (See also figure S7, S8, and S9).

### Structural implications of S31Δ and N30 glycosylation

As confirmed by mass spectrometry, S31Δ alters the “^30^NSFT^33^” sequence to “^30^N-FT^33^”, introducing an N30 glycosylated site on the NTD (**Figure 2A**). Structural alignment between JN.1 S (S31) and KP.3.1.1 S (S31Δ) in both “One RBD Up” and “All RBD Down” states revealed similar overall conformations (Cα RMSD = 1.1 Å and 0.9 Å, respectively) (**Figure 5A**) indicating that S31Δ and N30 glycosylation did not induce any substantial conformational changes. To observe a clear structure of the KP.3.1.1 NTD, we performed local refinement of the NTD and its adjacent RBD (**Figure 2B**). The map was of good quality and was resolved at 3.7 Å resolution (**Table S1 and Figure S2E**). Local refinement of KP.3.1.1 NTD showed that S31Δ results in a 180° flip of the F32 side chain rotamer into a hydrophobic pocket formed by L56, V62, Y91, F216, and Y266 (**Figure 5B**). Furthermore, the density of the NAG was clear at the N30 glycosylation site (**Figure 5B and 5C**). JN.1.11 showed a similar NTD structure to JN.1 (**Figure S8A and S8B**). Similarly, JN.1.11+S31Δ, JN.1.11.1+S31Δ, and JN.1.11+Q493E+S31Δ also shared similar NTD structures with KP.3.1.1 (**Figure S8C-S8F**). Three sarbecoviruses with N30 glycosylation (RaTG13, Pangolin-GD, and Pangolin-GX) also showed similar NTD structures and N30 glycosylation as KP.3.1.1 ^20–22^ (**Figure S8F-S8I**). In addition, N30 is located in NTD “site vi”, which comprises the binding sites for known antibodies, including neutralizing antibody C1717 and non-neutralizing antibodies DH1052, COV57, and S2M24 ^30–33^. Through structural comparisons, we show that N30 glycosylation is positioned such that it would likely affect the binding to mAbs C1717 and DH1052 (**Figure S9**).

### Analysis of the N-glycan landscape on the spike proteins

The coronavirus spike proteins are heavily glycosylated, and the SARS-CoV-2 wild-type spike protein has 22 N-glycosylation sites per protomer^34,35^. Glycan profiling studies by mass spectrometry have been performed for many early variants^36–38^, but here we also investigate the glycoforms present at three new glycosylation sites (N30, N245, N354). Compared to the WT spike protein, all of the seven spike proteins that we analyzed here exhibit two new glycosylation sites from wild-type (N245 in NTD and N354 in RBD) because of the H245N and K356T mutations that emerged in recent variants, while the N17 glycosylation in early variants is absent due to the T19I mutation that emerged in BA.2 and later variants ^27,39^. N245 is mostly occupied with complex glycans, while N354 is predominantly occupied with high mannose glycans. Notably, introduction of the S31Δ mutation in KP3.1.1 and engineered JN1.11/JN.1.11.1 spike variants created a glycosylation site at N30 that is mainly populated with high mannose glycans, reducing the percentage of complex glycans from ∼80-95% to <50% at the N61 glycosylation site (**Figure 2A**). Sequence alignment showed the N61 glycosylation site is highly conserved across all sarbecoviruses shown here (**Figure 2B**), suggesting a potentially critical, but not yet determined, role for the N61 glycan. A previous study has shown, however, that N61Q significantly reduces infectivity^40^. Zoomed-in views of KP.3.1.1, RaTG13, Pangolin-CoV-GD, and Pangolin-CoV-GX indicated that N30 and its attached glycan are spatially proximal to N61 (**Figure 2C**)^20–22^. We hypothesize that the shift of glycoforms at N61 is a consequence of crowding by the neighboring glycan at N30 and more restricted access to the glycan processing enzymes. Furthermore, α6-fucosylation of glycans at N61 would likely clash with N30 sugar moieties. Any potential impact of the altered N61 glycoform on its function remains to be investigated. Other N-glycans throughout the spike protein remain largely unchanged.

### Re-occurrence of residues in other sarbecoviruses during SARS-CoV-2 evolution

The Q493E mutation and additional N-glycosylation site at N30 resulting from S31Δ have only recently emerged in SARS-CoV-2 variants. This glycan has been observed in other clade-1b sarbecoviruses over a prolonged period (**Figure 6B**), such as pangolin-CoV GD, pangolin-CoV GX-P2V, and RaGT13^20–22^ (**Figure 6A**). The glycans in all four strains (including SARS-CoV-2 KP.3.1.1 and these three non-SARS-CoV-2 sarbecoviruses) exhibit similar conformations, implicating a conserved function of the glycan among sarbecoviruses (**Figures 2C and S8**). Furthermore, the Q493E mutation in SARS-CoV-2, which emerged in early 2024, is analogous to the E493 residue found in pangolin-CoV GX-P2V. In addition to Q493E and the gained N30 glycosylation site, other SARS-CoV-2 variants also exhibited re-occurrence of amino acids found in the spike proteins of other sarbecoviruses. In total, 15 mutations (S31Δ, S50L, F157S, L216F, R403K, F456L, N460K, T478K, V483del, F486P, Q493E, Y505H, P621S, H655Y, and D796Y) have been found in at least one SARS-CoV-2 variant of concern (VOC) or variant of interest (VOI) (**Figure 6B**). Notably, nine of these 15 re-occurring mutations (S31Δ, S50L, F157S, L216F, R403K, F456L, V483del, Q493E, and P621S), emerged only recently, suggesting an accelerated re-occurrence rate since late 2023. Data from previous deep mutational scanning (DMS) studies showed that most re-occurring mutations (R403K, F456L, F486P, Y505H, and P621S) performed better than average in hACE2 binding, cell entry, and RBD expression (**Figure 6D**)^16,41,42^. Although Q493E alone reduced hACE2 binding, it was complemented by F456L, which restored binding affinity. Overall, our analysis of major SARS-CoV-2 variants demonstrates that many of the predominant variants have reversions to amino acids originally found in other sarbecoviruses that may suggest alterations to the mutational landscape trajectory in recent years.

**Figure 6.**
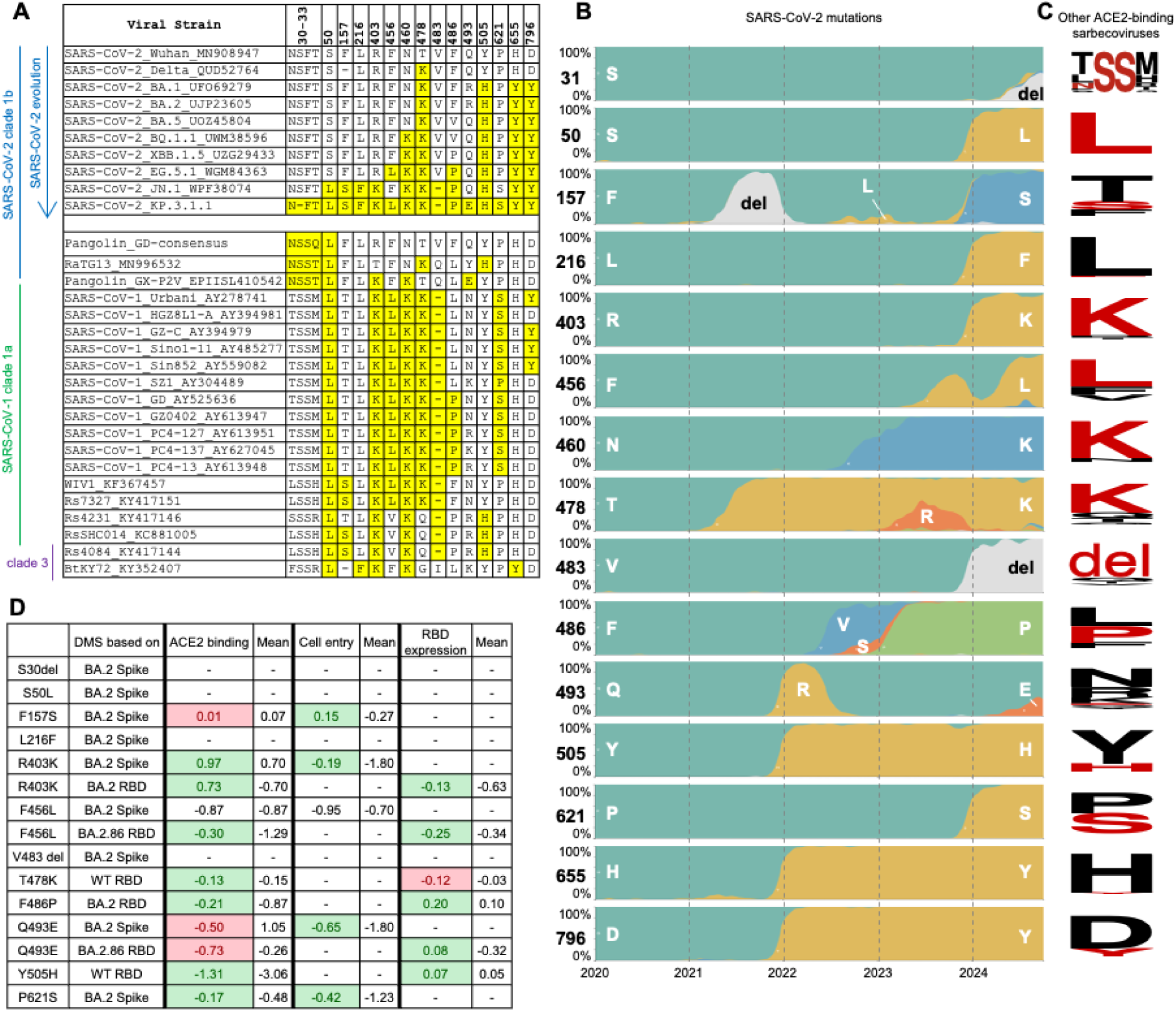
Re-occurrence of amino acids observed in other sarbecoviruses during SARS-CoV-2 evolution. **(A)** Sequence alignment shows recently mutated amino acids in SARS-CoV-2 that are found in other ACE2-binding sarbecoviruses^49^. Re-occurring amino acids are highlighted in yellow. Sequence accession codes are provided at the end of each viral strain name. **(B)** Frequency of amino acids at particular positions on SARS-CoV-2 spike as of October 2024. Data were sourced from nextstrain.org (GISAID data)^14,50^. **(C)** Sequence conservation analysis of ACE2-binding sarbecoviruses excluding SARS-CoV-2. Each row corresponds to the residue in panel B. Amino acids that have re-occurred in the currently dominant SARS-CoV-2 variant KP.3.1.1 are shown in red. The SARS-CoV-2 S31 deletion results in an N-glycosylation site ^30^NFT, corresponding to the N-glycosylation site ^30^NSS in some other sarbecoviruses, which is also highlighted here. The sequence conservation is plotted by WebLogo^51^. **(D)** Fold change in hACE2 binding affinity (Δlog10 K_D_), entry into 293T cells expressing high ACE2 (functional score), RBD expression (log10(MFI)) for assessment of re-occurring mutations. The average was calculated from all mutations at that site. Green highlights where the effect of the re-occurring mutation is greater than the average, and red highlights where the effect of the re-occurring mutation is less than the average. Data are from previous DMS studies^16,41,42^. (See also figure S10).

## Discussion

In this study, we used cryo-EM to determine high-resolution structures of the globally predominant variant KP.3.1.1 in both apo and ACE2-bound states. Our structural analysis of the KP.3.1.1 RBD/hACE2 complex and binding data are consistent with an epistatic interaction between the F456L and Q493E mutations in the RBD of KP.3.1.1, where Q493E alone substantially reduces binding but, in the presence of F456L, these residues synergize to restore ACE2 binding affinity. In the cryo-EM structure of the spike protein of KP.3.1.1, we observed N30 glycosylation and a 180° flip of the F32 sidechain, resulting from the S31 deletion in the ^30^NSFT^33^ motif. Additionally, we also determined apo spike protein structures of engineered JN.1-derived mutants carrying S31Δ, F456L, and Q493E. CryoSPARC 3D variability analysis demonstrated that all spike proteins had similar proportions of up and down RBDs, suggesting that S31Δ does not influence the conformational dynamics of the spike protein. Furthermore, S31Δ had minimal effect on ACE2 and BD55-1205 binding affinities and the overall spike thermostability. Glycan mass spectrometry demonstrates that glycans at the new N30 site are predominantly high mannose and alter the proportions of complex and high mannose/hybrid glycoforms at N61.

Previous functional studies indicated that, compared to KP.3, KP.3.1.1 exhibits greater resistance to neutralizing antibodies from convalescent plasma, immunized rabbit sera, and vaccinated human sera, as well as higher infectivity and reduced binding affinity to monoclonal antibodies across multiple classes^8,10,11,15,28^. Additionally, the recently emerged XEC variant, which has another N-glycosylation in the NTD due to a T22N mutation, but does not have the S31 deletion or N30 glycan (**Figure S10**), appears to display a similar extent of immune evasion as KP.3.1.1^10,15,28^. Although we do not have sufficient evidence to link the addition of NTD glycans to immune invasion based only on the current structural analysis of recombinant spike proteins, we hypothesize that S31Δ-introduced N30 glycosylation contributes to immune evasion of KP.3.1.1. Further functional studies in live viruses are needed to elucidate the underlying mechanism.

KP.3 contains the F456L and Q493E mutations, which exhibit synergistic effects, significantly enhancing ACE2 binding affinity compared to Q493E mutation alone^16,17^. In this study, we demonstrate that KP.3.1.1 retains this synergistic epistasis. Epistasis is a common phenomenon in RNA viruses, including vesicular stomatitis virus (VSV)^43^, HIV-1^44^, and influenza virus^45,46^. It has been shown to play a vital role in enhancing viral fitness and enabling immune escape in SARS-CoV-2^42,47^. Investigating the effects of mutations in different SARS-CoV-2 variants and other sarbecoviruses may help reveal epistatic interactions that may confer fitness advantages and shape the evolutionary trajectories of SARS-CoV-2.

The evolution trajectory of SARS-CoV-2 is largely constrained and driven by immune pressure while maintaining or enhancing the viral fitness. The immunity of the global population has been largely imprinted by early SARS-CoV-2 strains, either through vaccination or early infections. While SARS-CoV-2 originally diverged from related sarbecoviruses, many amino acids were first seen and unique to the SARS-CoV-2. Mutations at these residues have facilitated immune escape but sometimes at the cost of viral fitness. Interestingly, amino acids at similar positions in non-SARS-CoV-2 sarbecoviruses, which have demonstrated high viral fitness, are now re-emerging in SARS-CoV-2. These re-occurring mutations likely represent a balance between immune escape and fitness optimization. Notably, we identified 15 mutations during SARS-CoV-2 evolution that correspond to amino acids present in other sarbecoviruses. These findings highlight the evolutionary trajectories that are now possible and productive and are bringing the spike proteins in these sarbecoviruses closer together. These re-occurring mutations now appear to confer some structural and functional advantages as SARS-CoV-2 continues to evolve to maintain fitness and immune escape, offering information on possible evolutionary trajectories for the next set of SARS-CoV-2 variants. Overall, the amino acid re-occurrence in related sarbecoviruses, along with the epistatic mutations observed during SARS-CoV-2 evolution, play critical roles in enhancing viral fitness and facilitating immune escape. These observations provide invaluable insights for predicting future evolutionary pathways of SARS-CoV-2.

## Acknowledgments

We thank Hannah Turner and Anant Gharpure for technical support with the electron microscopy, Charles Bowman and Marc-André Elsliger for assistance with computation, Jeanne Matteson and Beverly Ellis for contribution to mammalian cell culture, and Jeffrey Copps for protein purification. This study was supported by the Bill and Melinda Gates Foundation INV-004923 (I.A.W. and A.B.W.) and National Institute of Allergy and Infectious Diseases UM1 AI144462 (Scripps Consortium for HIV/AIDS Vaccine Development; J.C.P and J.R.Y).

## Author contributions

Z.F., I.A.W., and A.B.W. conceived the study. I.A.W. and A.B.W. supervised the study. Z.F. performed cryo-EM data collection. Z.F., J.H., and S.B. (Sandhya Bangaru) processed the cryo-EM data and built the model. Z.F. performed binding assays and interpreted data. S.B. (Sabyasachi Baboo) and J.K.D. performed mass spectrometry-based glycan analysis and data interpretation. J.C.P. and J.R.Y. supervised glycan analyses. M.Y. performed the bioinformatic study. Z.F., M.Y., I.A.W, and A.B.W. wrote the paper, and all authors reviewed and edited the paper. The funding was secured by J.C.P., J.R.Y., I.A.W., and A.B.W.

## MATERIALS AND METHODS

### Expression and purification of human ACE2

For cryo-EM and biolayer interferometry (BLI) binding experiments, human ACE2 (residues 19-615, GenBank: BAJ21180.1) was flanked with an N-terminal signal peptide and a C-terminal His_6_ tag and codon optimized in phCMV3 expression vector. Expi293S cells (Life Technologies) were then transfected with ExpiFectamine 293. Reagent and incubated according to manufacturer’s instruction. Supernatants were harvested and proteins were purified by Ni Sepharose excel resin (Cytiva), eluted with 300 mM imidazole, followed by size exclusion (SEC) and buffer exchanged into 20 mM Tris-HCl, 150 mM NaCl, pH 7.4.

### Expression and purification of BD55-1205 Fab

The heavy and light chains of the Fab were cloned into the phCMV3 vector. The plasmids were transiently co-transfected into ExpiCHO cells at a ratio of 2:1 (heavy chain to light chain) using ExpiFectamine CHO Reagent (Thermo Fisher Scientific) according to the manufacturer’s instructions. The supernatant was collected at 7 days post-transfection. The Fab was purified with a CaptureSelect CH1-XL Pre-packed Column (Thermo Fisher Scientific), followed by SEC and buffer exchanged 20 mM Tris-HCl, 150 mM NaCl, pH 7.4.

### Recombinant full-length SARS-CoV-2 prefusion spike protein and mutants

All spike ectodomain constructs contain a C-terminal T4 fibritin trimerization domain^52^ and a Twin-strep-tag for purification. For the SARS-CoV-2 spike, we synthesized a base construct (HP-GSAS) with residues 1 to 1208 from the Wuhan-Hu-1 strain (GenBank: QHD43416.1) with six stabilizing proline (HexaPro or S6P)^53^ substitutions at positions 817, 892, 899, 942, 986, and 987 and the S1/S2 furin cleavage site modified to 682-GSAS-685. The S6P expression construct containing the Omicron BA.4/BA.5 mutations (T19I, L24S, del25-27, del69-70, G142D, V213G, G339D, S371F, S373P, S375F, T376A, D405N, R408S, K417N, N440K, G446S, L452R, S477N, T478K, E484A, F486V, Q498R, N501Y, Y505H, D614G, H655Y, N658S, N679K, P681H, N764K, D796Y, Q954H, N969K) were assembled from synthesized DNA fragments. XBB.1.5, BA.2.86, JN.1, JN.1.11, JN.1.11.1, KP.3.1.1, and other mutants (JN.1.11+S31Δ, JN.1.11+Q493E, JN.1.11+S31Δ+Q493E, JN.1.11.1+S31Δ, and KP.3.1.1) were also synthesized by this method.

### Expression and purification of recombinant spike proteins

For protein expression, Expi293F cells (Thermo Fisher Scientific: A14527, RRID: CVCL_D615) and FreeStyle 293-F cells (Thermo Fisher Scientific: R79007, RRID: CVCL_D603) were transfected with the spike plasmid of interest. Transfected cells were incubated at 37°C with shaking at 110 r.p.m. for 6 days for protein expression. The spike proteins were purified from the supernatants with Strep-Tactin^®^XT 4Flow^®^ high capacity (IBA Lifescience GmbH: 2-5010-025) and buffer-exchanged to TBS (20 mM tris and 150 mM NaCl, pH 7.4) before further purification with SEC. Protein fractions corresponding to the trimeric spike proteins were collected and concentrated.

### Biolayer interferometry (BLI) binding assays

Binding assays were performed by biolayer interferometry (BLI) using an Octet Red instrument (FortéBio). Twin-strep-tagged spike proteins at 20 μg/mL in 1x kinetics buffer (1x PBS, pH 7.4, 0.01% BSA and 0.002% Tween 20) were loaded onto streptavidin (SA) sensors, and then incubated with 12.5, 25, 50, 100, 200, and 400 nM of hACE2 or 8.33, 25, and 75 nM of BD55-1205 Fab. The assay consisted of five steps: 1) baseline: 60 s with 1x kinetics buffer; 2) loading: 180s with Twin-strep-tagged spike proteins; 3) baseline: 90 s with 1x kinetics buffer; 4) association: 240s with hACE2 or antibody; and 5) dissociation: 240s with 1x kinetics buffer. For estimating the K_D_ values, a 1:1 binding model was used.

### Cryo-EM sample preparation

For JN.1.11, JN.1.11+S31Δ, KP.3.1.1, 3 µL of spike protein in TBS (20 mM Tris and 150 mM NaCl, pH 7.4) at 0.7 mg/mL was mixed with 0.5 µL of 0.035 mM lauryl maltose neopentyl glycol (LMNG) solution immediately before sample deposition onto an UltrAuFoil 1.2/1.3 (Au, 300-mesh; Quantifoil Micro Tools GmbH) grid that had been plasma cleaned for 25 seconds using a PELCO easiGlow™ Glow Discharge Cleaning System (Ted Pella Inc.). Following sample application, grids were blotted for 3.5 seconds before being vitrified in liquid ethane using a Vitrobot Mark IV (Thermo Fisher). For JN.1.11.1+S31Δ and JN.1.11+S31Δ+Q493E, blot time was 3 seconds. The KP.3.1.1 spike protein was incubated with a 3-fold molar excess of ACE2 (protomer: ACE2 = 1:1, final concentration of 0.75 mg/ml) for 30 min and following the same procedure as for the apo samples with 3 seconds blot time. The grids with glassy ice were stored in liquid nitrogen before data collection.

### Cryo-EM data collection and processing

Data collection was performed on a Thermo Fisher Glacios operating at 200 keV mounted with a Thermo Fisher Falcon 4 direct electron detector using the Thermo Fisher EPU 2 software at a magnification of 190,000x, resulting in a 0.718 Å pixel size. All micrographs were collected at a total cumulative dose of 44.84 e^-^/Å^2^. CryoSPARC Live Patch Motion Correction was used for alignment and dose weighting of movies. CTF estimations, particle picking, particle extraction, iterative rounds of 2D classification, ab-Initio reconstruction, heterogeneous refinement, homogeneous refinement, 3D Variability, global CTF refinement, and non-uniform refinement were performed on CryoSPARC^54^.

### Model building and refinement

Initial model building was performed manually in Coot using JN.1 “One RBD Up” S (PDB ID: 8Y5J), JN.1 “All RBD Down” S (PDB ID: 8X4H), and JN.1 S/hACE2 complex (PDB ID: 8Y16) as the templates^1,27^. Followed by iterative rounds of Rosetta relaxed refinement, Phenix real space refinement, and Coot manual refinement to generate the final models^55–57^. EMRinger and MolProbity were run following each round of Rosetta refinement to evaluate and choose the best refined models^58,59^. Final map and model statistics are summarized in **Table S1**. Figures were generated using PyMOL, UCSF Chimera, and UCSF Chimera X^60,61^.

### Nano Differential Scanning Fluorimetry

NanoDSF was performed using Prometheus PANTA (NanoTemper Technologies, München, Germany). Samples were loaded in nanoDSF grade standard capillaries (NanoTemper Technologies GmbH, München, Germany) and exposed at thermal stress from 20°C to 95°C by thermal ramping rate of 1°C/min. Fluorescence emission from tryptophan after UV excitation at 280 nm was collected at 330 nm and 350 nm with a dual-UV detector. Thermal stability parameters *T*_m_ was calculated by Panta Analysis software (NanoTemper Technologies, München, Germany).

### Mass spectrometry-based N-glycan analysis

DeGlyPHER ^23,62^ was used to ascertain site-specific glycan occupancy and processivity on the examined glycoproteins.

### Proteinase K and trypsin treatment

Recombinant SARS-CoV-2 S-glycoprotein was exchanged to water using the Microcon Ultracel PL-10 centrifugal filter. Glycoprotein was reduced with 5 mM tris (2-carboxyethyl) phosphine hydrochloride (TCEP-HCl) and alkylated with 10 mM 2-Chloroacetamide in 100 mM ammonium acetate for 20 min at room temperature (RT, 24°C). Glycoprotein was digested with 1:25 Proteinase K (PK) for 30 min at 37°C and subsequently analyzed by LC-MS/MS.

### LC-MS/MS

Samples were analyzed on an Orbitrap Eclipse Tribrid mass spectrometer. Samples were injected directly onto a 25 cm, 100 μm ID column packed with BEH 1.7 μm C18 resin. Samples were separated at a flow rate of 300 nL/min on an EASY-nLC 1200 UHPLC. Buffers A and B were 0.1% formic acid in 5% and 80% acetonitrile, respectively. The following gradient was used: 1–25% B over 100 min, an increase to 40% B over 20 min, an increase to 90% B over another 10 min and held for 10 min at 90% B for a total run time of 140 min. The column was re-equilibrated with buffer A prior to the injection of the sample. Peptides were eluted from the tip of the column and nanosprayed directly into the mass spectrometer by application of 2.8 kV at the back of the column. The mass spectrometer was operated in a data-dependent mode. Full MS1 scans were collected in the Orbitrap at 120,000 resolution. The cycle time was set to 3s and, within this 3s, the most abundant ions per scan were selected for HCD MS/MS at 35 NCE. Dynamic exclusion was enabled with exclusion duration of 60 seconds and singly charged ions were excluded.

### LC-MS/MS Data Processing

Protein and peptide identification were carried out with the Integrated Proteomics Pipeline (IP2). Tandem mass spectra were extracted from raw files using RawConverter ^63^ and searched with ProLuCID ^64^ against a database comprising UniProt reviewed (Swiss-Prot) proteome for *Homo sapiens* (UP000005640), UniProt amino acid sequences for Endo H (P04067), PNGase F (Q9XBM8), and Proteinase K (P06873), amino acid sequences for the examined proteins, and a list of general protein contaminants. The search space included no cleavage specificity. Carbamidomethylation (+57.021460 C) was considered a static modification. Deamidation in the presence of H_2_ ^18^O (+2.988261 N), GlcNAc (+203.079373 N), oxidation (+15.994915 M), and N-terminal pyroglutamate formation (–17.026549 Q) were considered differential modifications. Data were searched with 50 ppm precursor ion tolerance and 50 ppm fragment ion tolerance. We utilized a target-decoy database search strategy to limit the false discovery rate to 1%, at the spectrum level ^65^. A minimum of one peptide per protein, and no tryptic end per peptide for PK and one tryptic end per peptide for trypsin were required and precursor delta mass cut-off was fixed at 10 ppm. Statistical models for peptide mass modification (modstat) were applied. Census2 ^66^ label-free analysis was performed based on the precursor peak area, with a 10 ppm precursor mass tolerance and 0.1 min retention time tolerance. Data analysis using GlycoMSQuant ^23^ was implemented to automate the analysis. GlycoMSQuant summed precursor peak areas across replicates, discarded peptides without NGS, discarded misidentified peptides when N-glycan remnant-mass modifications were localized to non-PNGS asparagines, and corrected/fixed N-glycan mislocalization where appropriate. Resultant PNGS were aligned to the SARS-CoV-2 (Wuhan-Hu-1) S-protein sequence from UniProt (P0DTC2).

