## Supplemental Material for "Structural and Functional Insights into the Evolution of SARS-CoV-2 KP.3.1.1 Spike Protein"

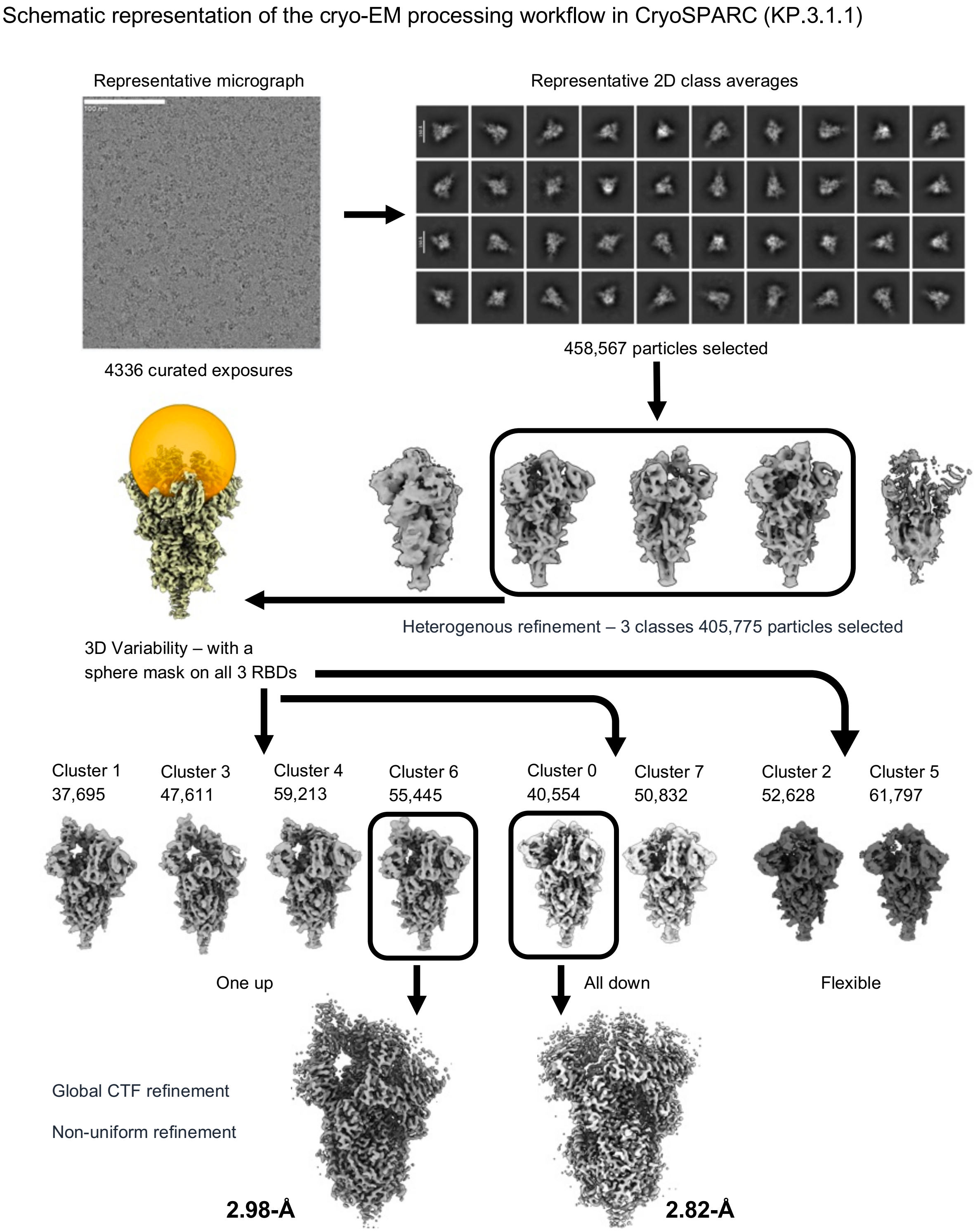


**Figure S1. Schematic representation of the cryo-EM processing workflow for SARS-CoV-2 KP.3.1.1 apo spike.** Workflow in cryoSPARC including initial processing steps (motion correction, CTF estimation, micrograph selection, particle picking, and selection based on 2D classification and heterogenous refinement) and 3D variability with a sphere mask on all three RBDs. All particles were then separated into eight clusters and classified by their conformation. A representative map from each class underwent global CTF refinement and non-uniform refinement and was used for model building. This workflow has been applied to all datasets.

**A**

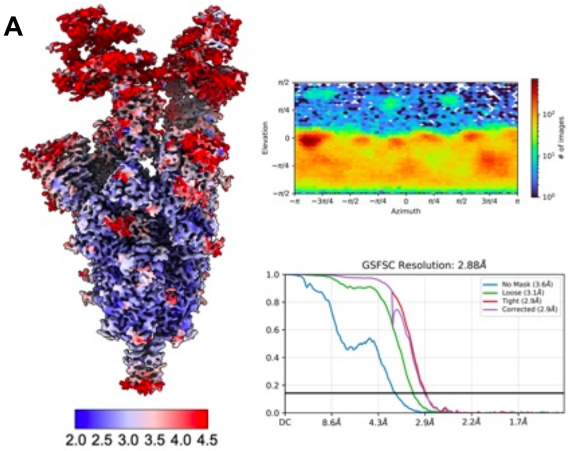


KP.3.1.1 S + hACE2

**B**

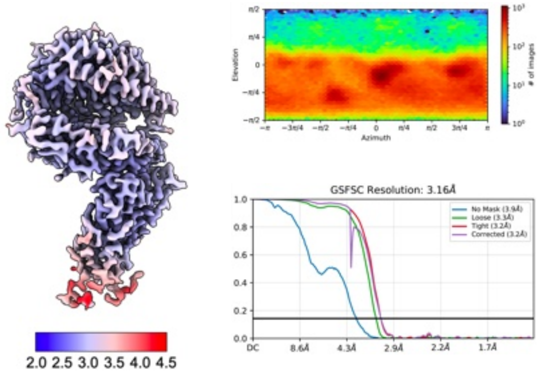


KP.3.1.1 RBD + hACE2

**C**

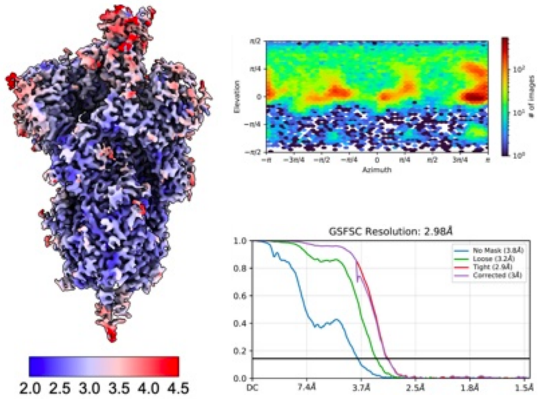


KP.3.1.1 S "One RBD Up"

**D**

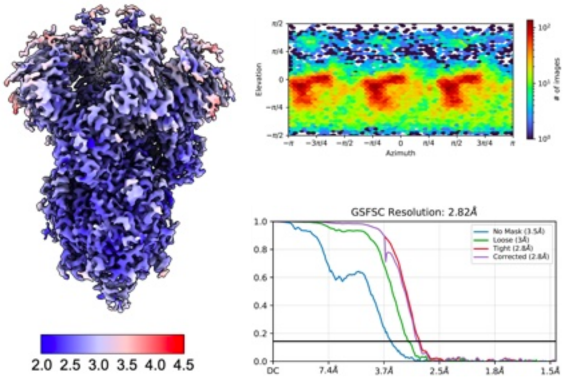


KP.3.1.1 S "All RBD Down"

**E**

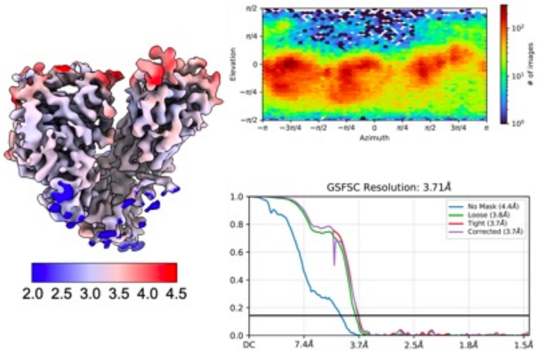


KP.3.1.1 RBD + NTD

F

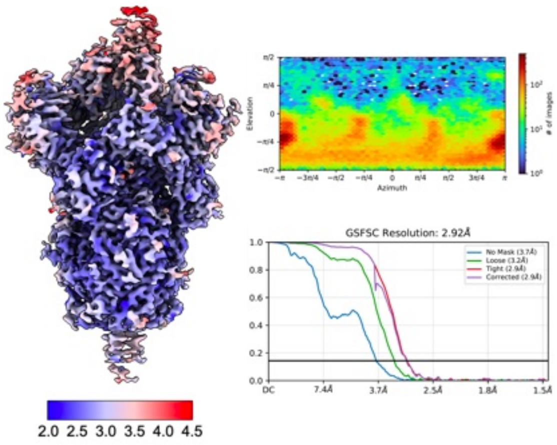


JN.1.11+S31Δ+Q493E S “One RBD Up”

G

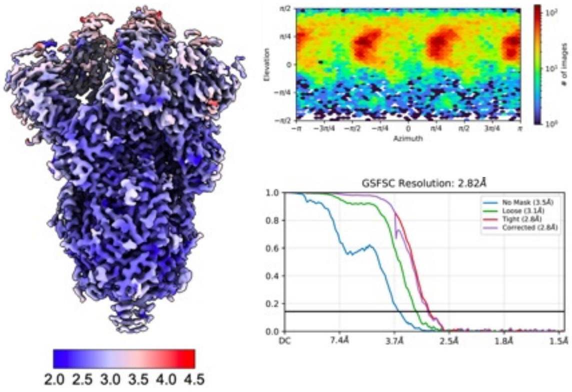


JN.1.11+S31Δ+Q493E S “All RBD Down”

H

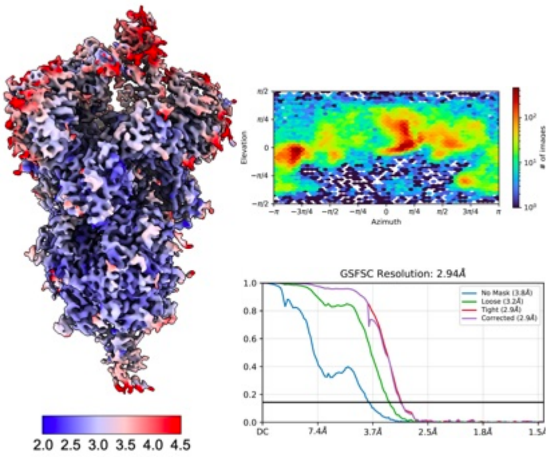


JN.1.11.1+S31Δ S “One RBD Up”

I

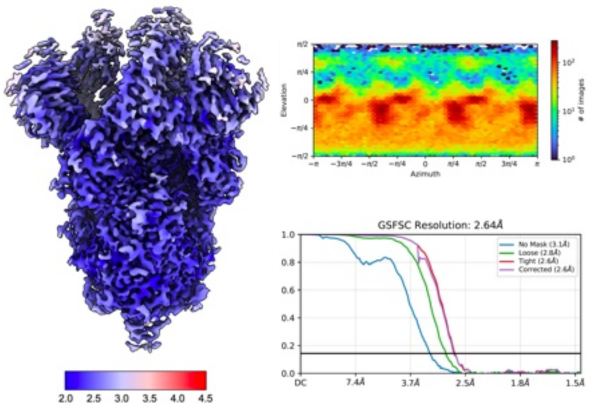


JN.1.11.1+S31 Δ S “All RBD Down”

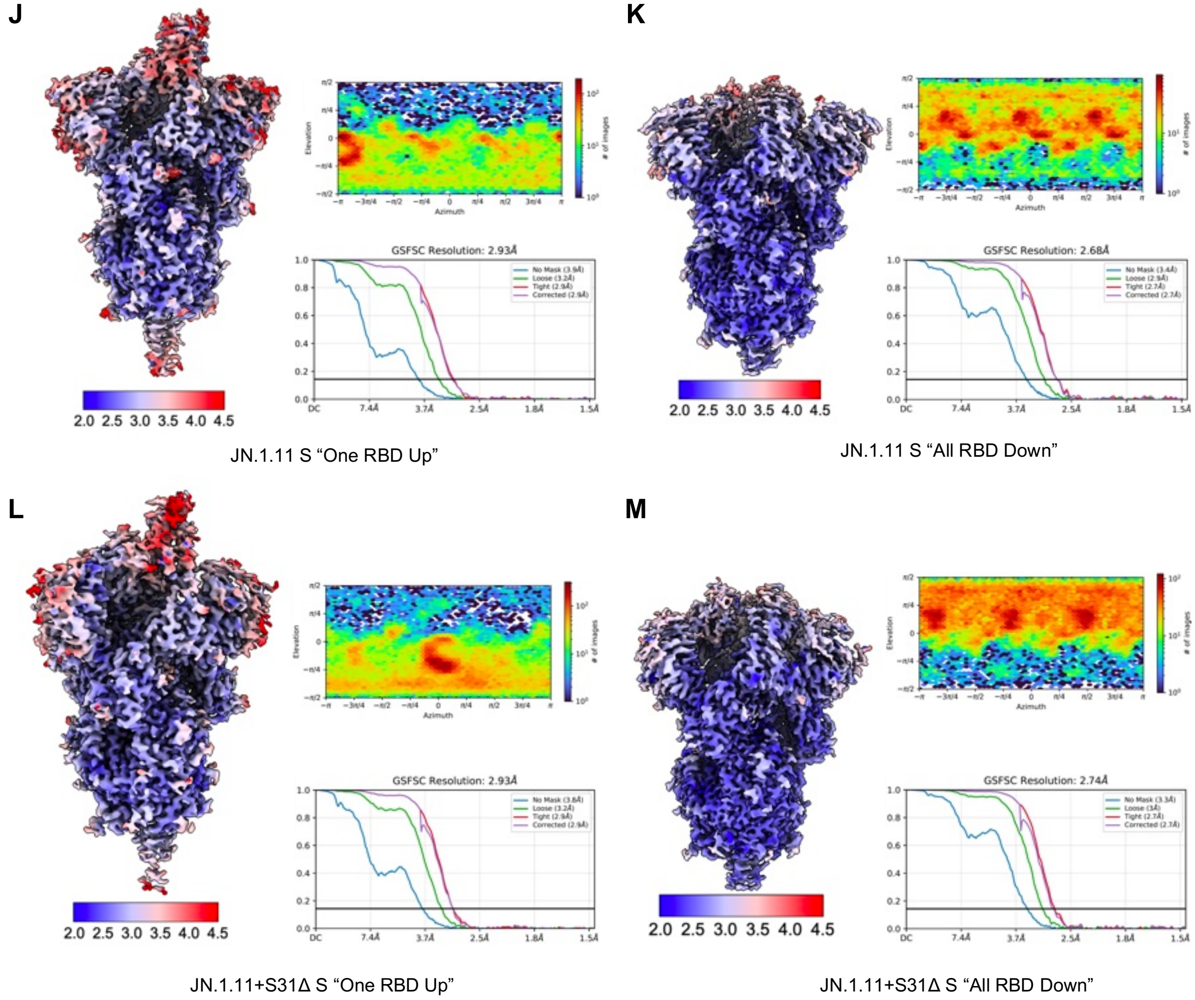


**Figure S2. Angular distribution, corresponding FSC curves, and resolution distribution of the cryo-EM map.** In the global and local resolution distribution of the cryo-EM maps, blue represents the high-resolution areas (2 Å) while red represents the low-resolution (4.5Å). **(A)** KP.3.1.1 S + hACE2. **(B)** KP.3.1.1 RBD + hACE2. **(C)** KP.3.1.1 S "One RBD Up". **(D)** KP.3.1.1 S "All RBD Down". **(E)** KP.3.1.1 RBD + NTD. **(F)** JN.1.11+S31Δ+Q493E S "One RBD Up". **(G)** JN.1.11+S31Δ+Q493E S "All RBD Down". **(H)** JN.1.11.1+S31Δ S "One RBD Up". **(I)** JN.1.11.1+S31 Δ S "All RBD Down". **(J)** JN.1.11 S "One RBD Up". **(K)** JN.1.11 S "All RBD Down". **(L)** JN.1.11+S31Δ S "One RBD Up". **(M)** JN.1.11+S31Δ S "All RBD Down".

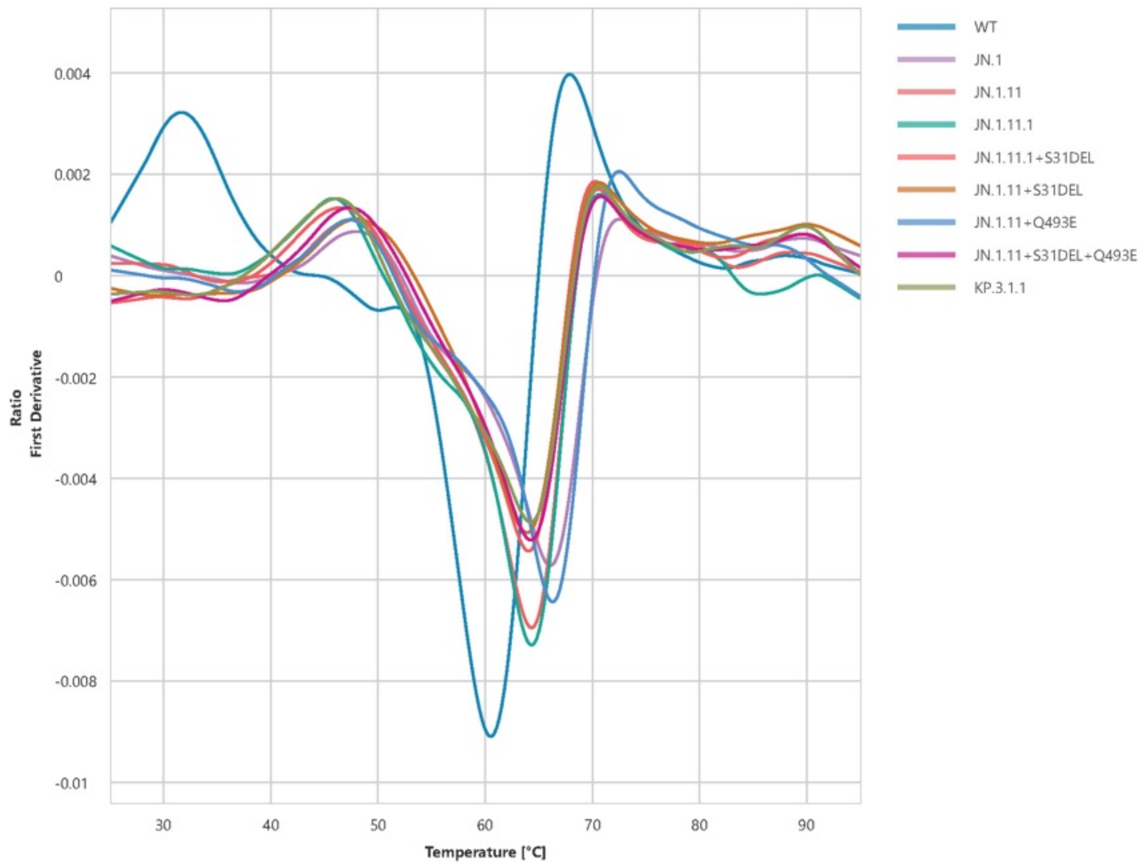


| Sample | Tm (°C) |
|---|---|
| WT | 31.65 |
| JN.1 | 47.87 |
| JN.1.11 | 47.59 |
| JN.1.11.1 | 45.71 |
| JN.1.11.1+S31DEL | 46.46 |
| JN.1.11+S31DEL | 48.03 |
| JN.1.11+Q493E | 47.29 |
| JN.1.11+S31DEL+Q493E | 47.31 |
| KP.3.1.1 | 46.01 |

**Figure S3. Differential scanning fluorimetry analysis of spike variant thermostability.**

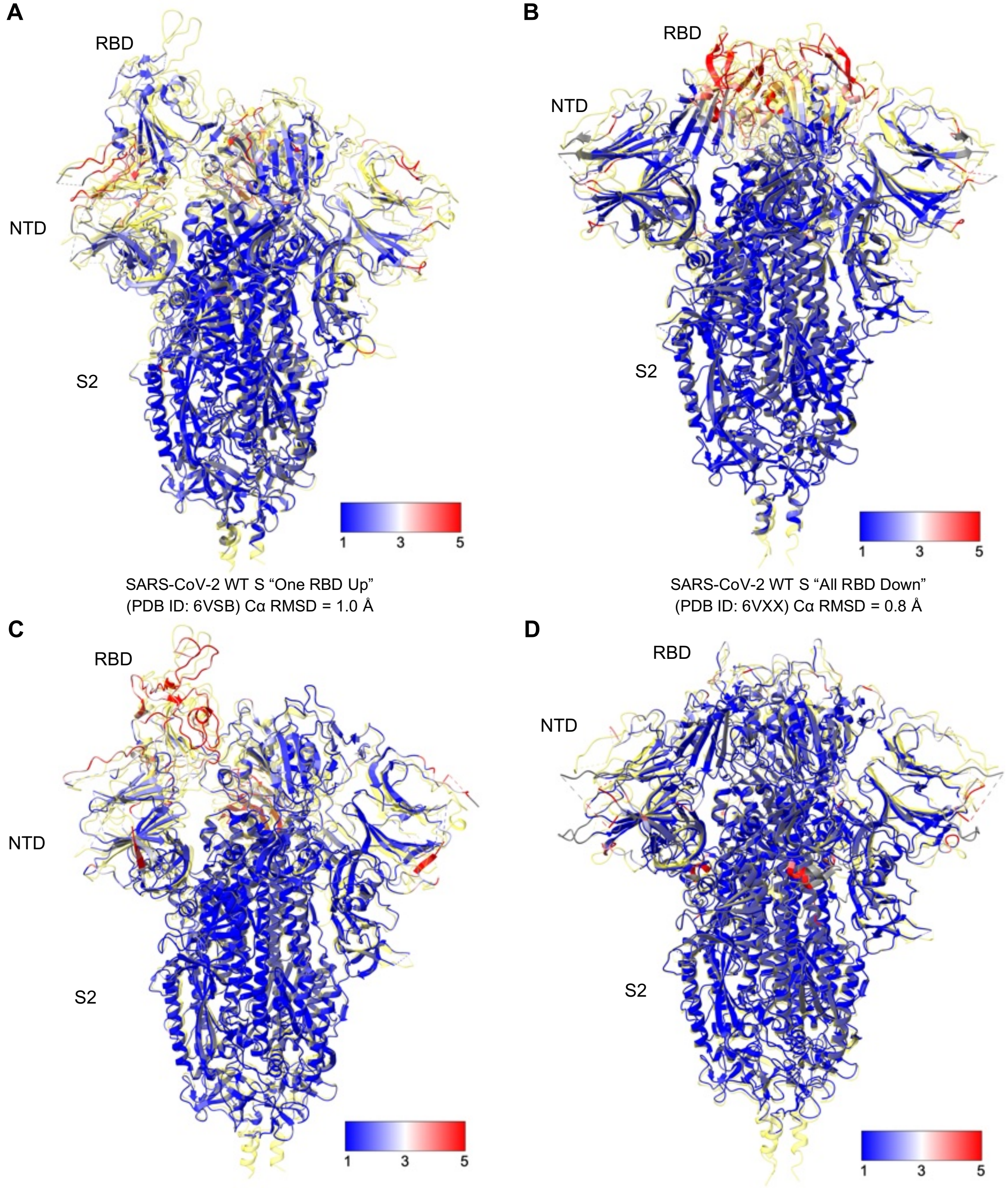


SARS-CoV-2 WT S “One RBD Up”
(PDB ID: 6VSB) Cα RMSD = 1.0 Å

SARS-CoV-2 WT S “All RBD Down”
(PDB ID: 6VXX) Cα RMSD = 0.8 Å

SARS-CoV-2 JN.1 S “One RBD Up”
(PDB ID: 8Y5J) Cα RMSD = 1.1 Å

SARS-CoV-2 JN.1 S “All RBD Down”
(PDB ID: 8X4H) Cα RMSD = 0.9 Å

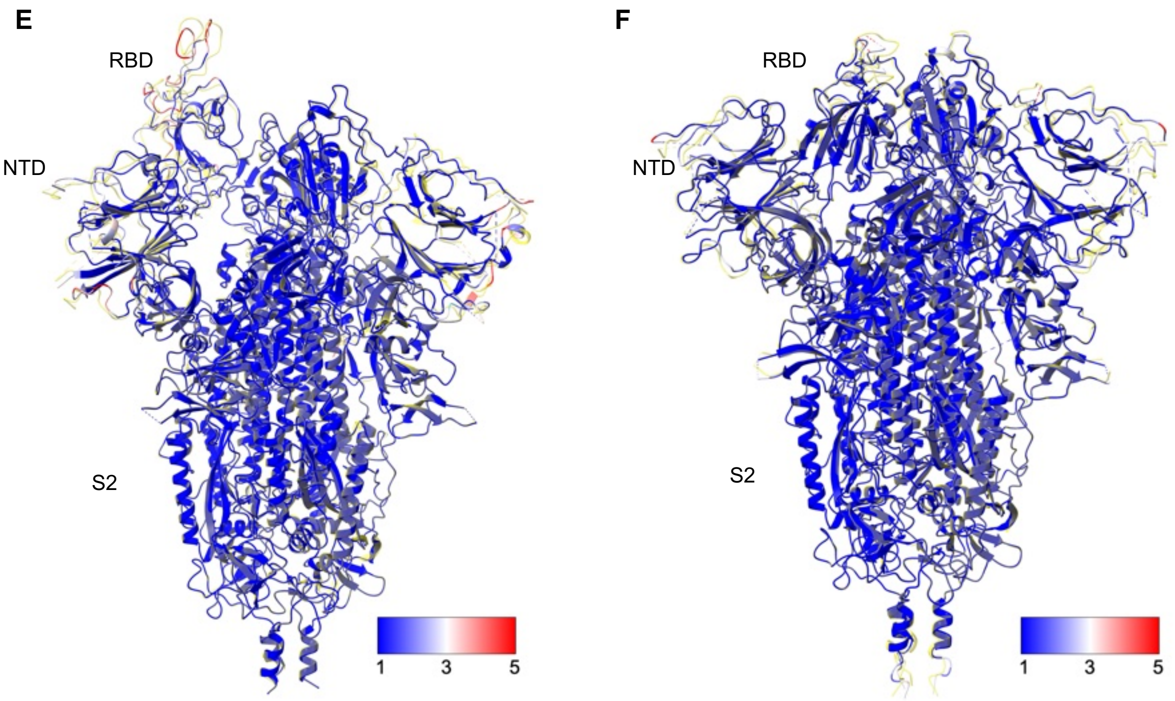


SARS-CoV-2 JN.1.11 S "One RBD Up"
(This study) Cα RMSD = 0.6 Å

SARS-CoV-2 JN.1.11 S "All RBD Down"
(This study) Cα RMSD = 0.5 Å

**Figure S4. Structural superimposition of SARS-CoV-2 WT, JN.1, and JN.1.11 spike protein with KP.3.1.1 spike protein.** KP.3.1.1 spike proteins are colored light yellow. Structural differences of SARS-CoV-2 WT, JN.1, and JN.1.11 spike are color-coded by their root mean square deviation (RMSD) (Å). Positions of RBD, NTD, and S2 are labeled. **(A, C, E)** Structural alignment of the KP.3.1.1 "One RBD Up" spike structure to SARS-CoV-2 WT (PDB ID: 6VSB), JN.1 (PDB ID: 8Y5J), and JN.1.11 (this study), respectively. **(B, D, F)** Structural alignment of the KP.3.1.1 "All RBD Down" spike structure to SARS-CoV-2 WT (PDB ID: 6VXX), JN.1 (PDB ID: 8X4H), and JN.1.11 (this study), respectively.

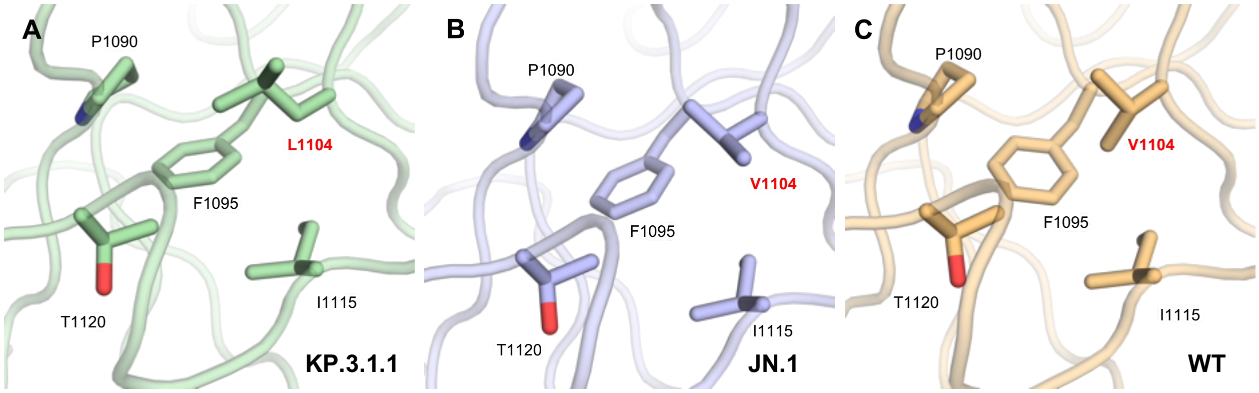


**Figure S5. Zoomed-in view of V/L1104 in WT, JN.1, and KP.3.1.1 spikes. (A-C)** L1104 or V1104 is surrounded by a hydrophobic pocket formed by P1090, F1095, I1115, and T1120 in KP3.1.1., JN.1, and WT spike proteins.

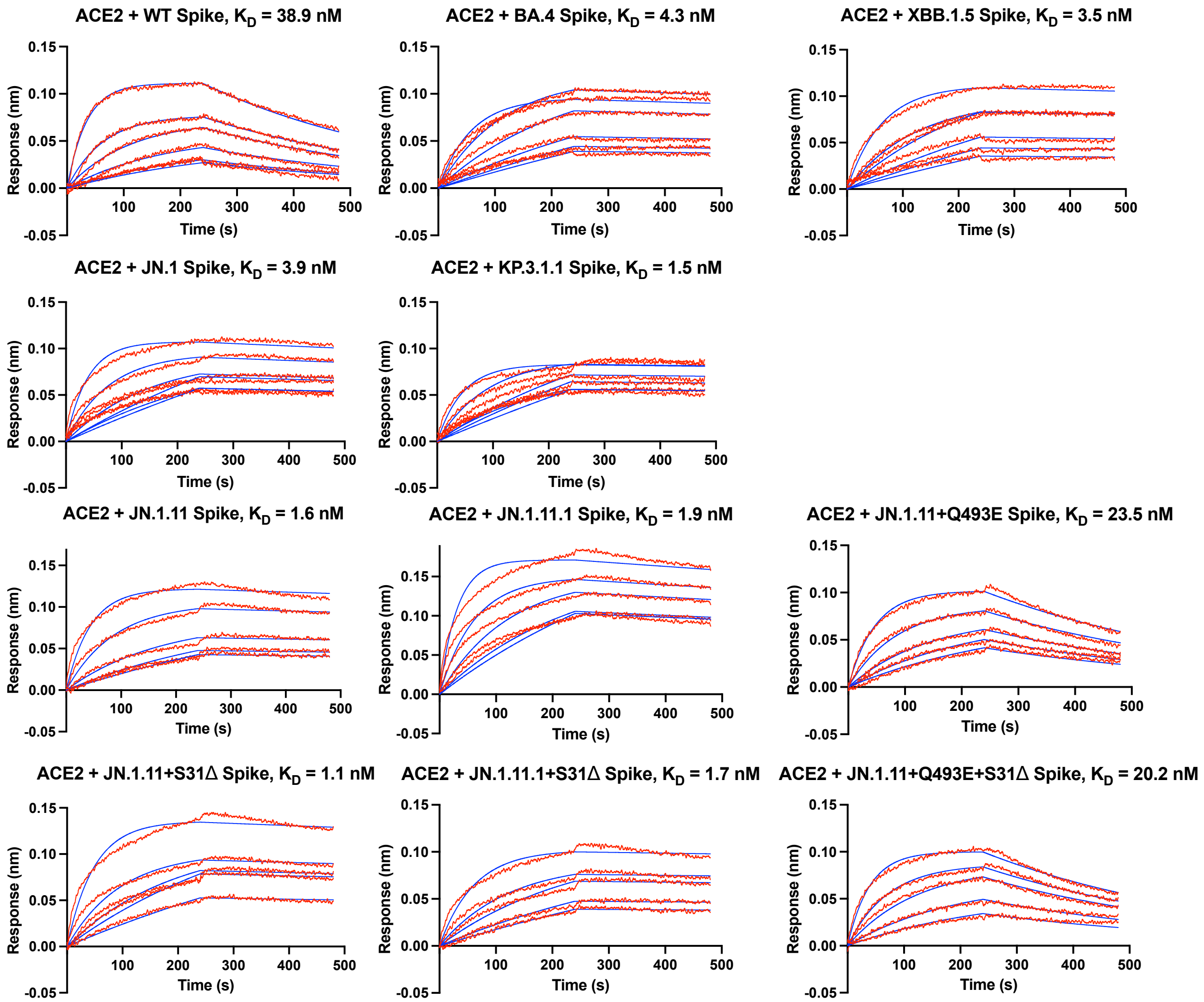


**Figure S6. Sensorgrams for binding of various spikes to hACE2, related to Fig. 4F.** Binding kinetics of different spike proteins against hACE2 were measured by biolayer interferometry (BLI). The Y-axis represents the response. Red lines represent the response curve, and blue lines represent a 1:1 binding model. Binding kinetics were measured for hACE2 concentrations (12.5, 25, 50, 100, 200, and 400 nM). Dissociation constant ($K_D$) is indicated.

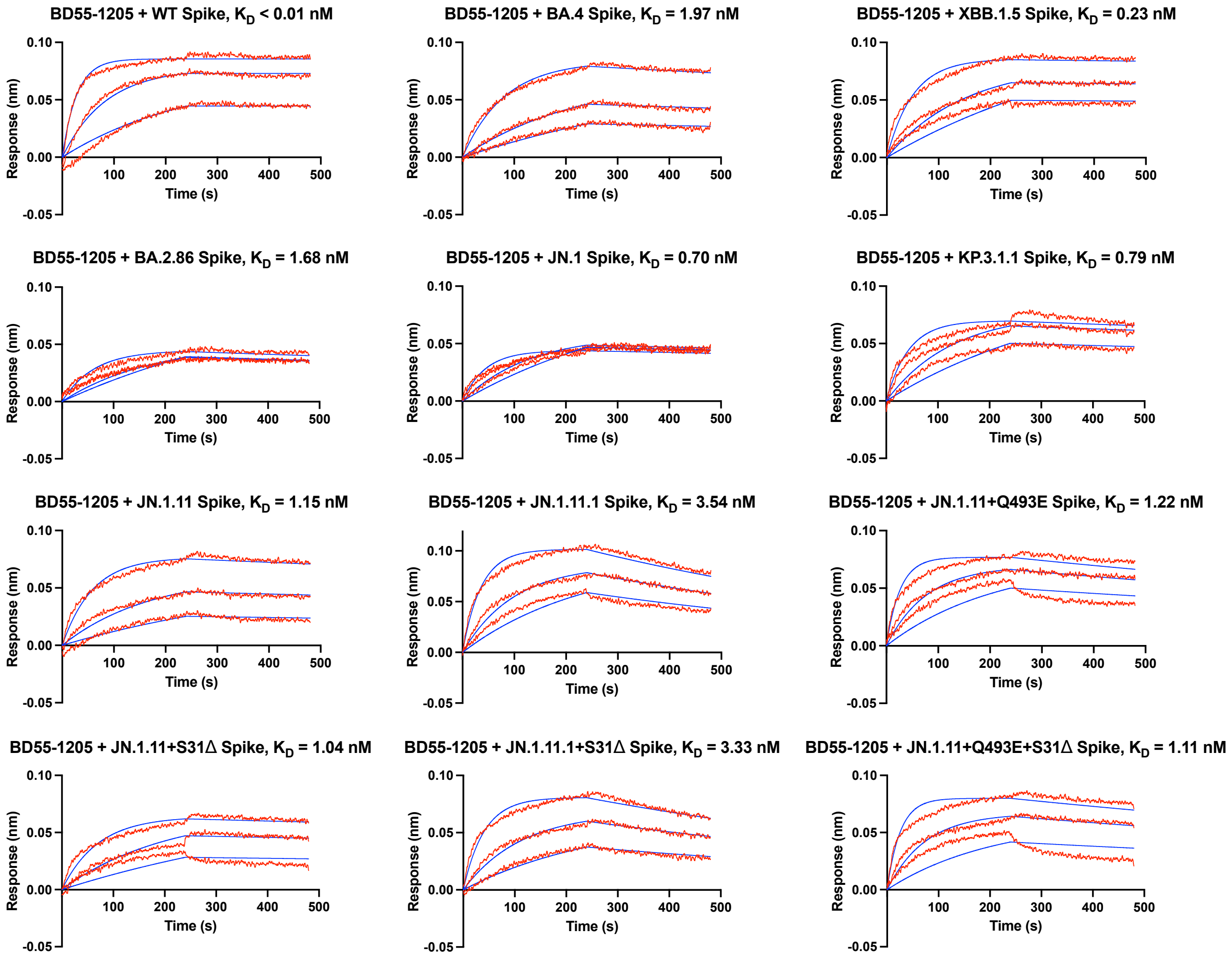


**Figure S7. Sensorgrams for binding of various spikes to BD55-1205 Fab, related to Fig. 5F.** Binding kinetics of different spike proteins against BD55-1205 Fab were measured by biolayer interferometry (BLI). The Y-axis represents the response. Red lines represent the response curve, and blue lines represent a 1:1 binding model. Binding kinetics were measured for Fab concentrations (8.33, 25, and 75 nM). Dissociation constant ($K_D$) is indicated.

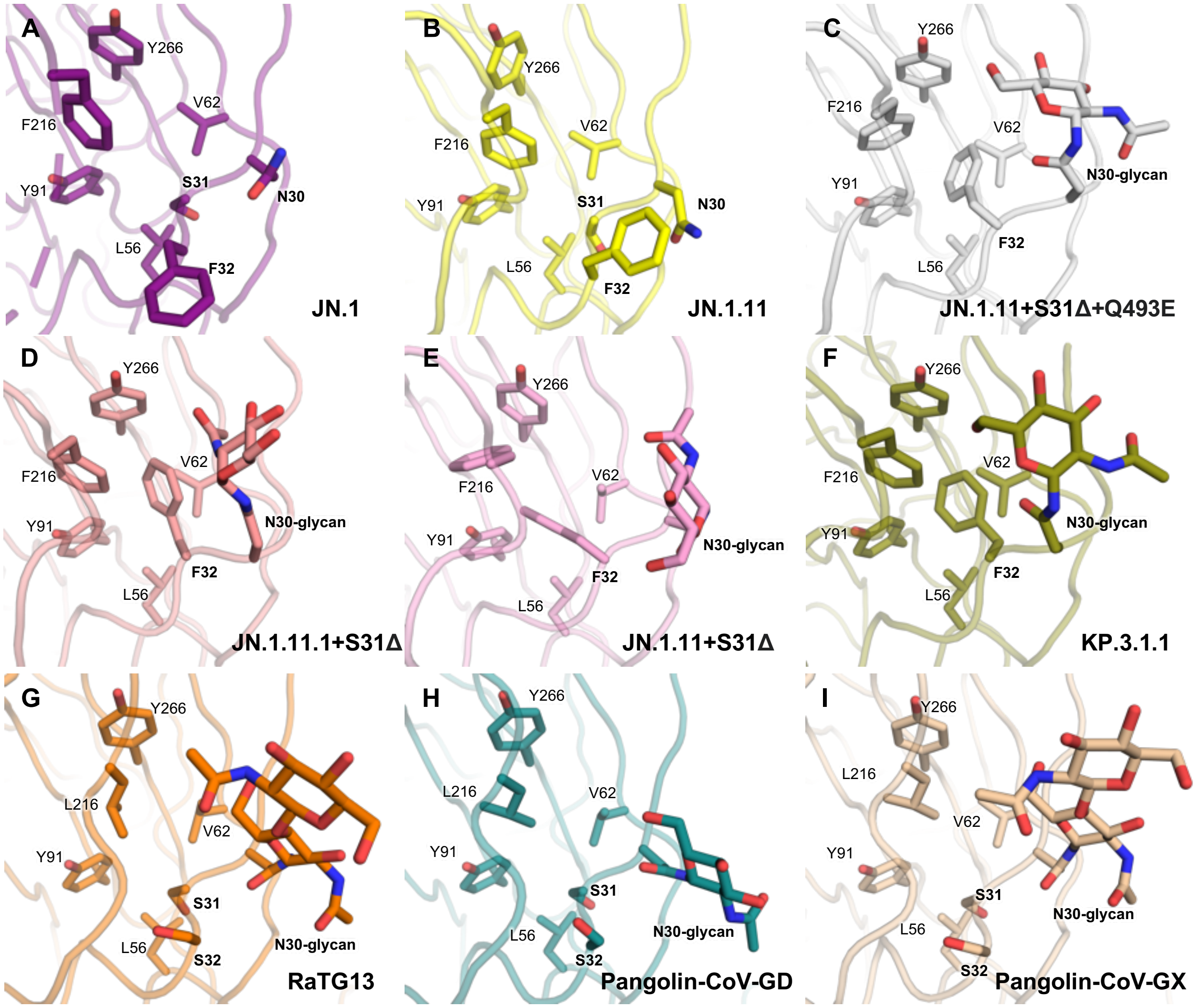


**Figure S8. Zoomed-in view of N30 in various spikes. (A-I)** Detailed side chain conformation including N30 with glycan, S31, F/S32, L56, V62, Y91, L/F216, and Y266 of JN.1 (purple), JN.1.11 (yellow), JN.1.11+S31Δ+Q493E (grey), JN.1.11.1+S31Δ (flesh), JN.1.11+S31Δ (pink), KP.3.1.1 (olive), RaTG13 (orange), Pangolin-CoV-GD (teal), and Pangolin-CoV-GX (wheat), respectively.

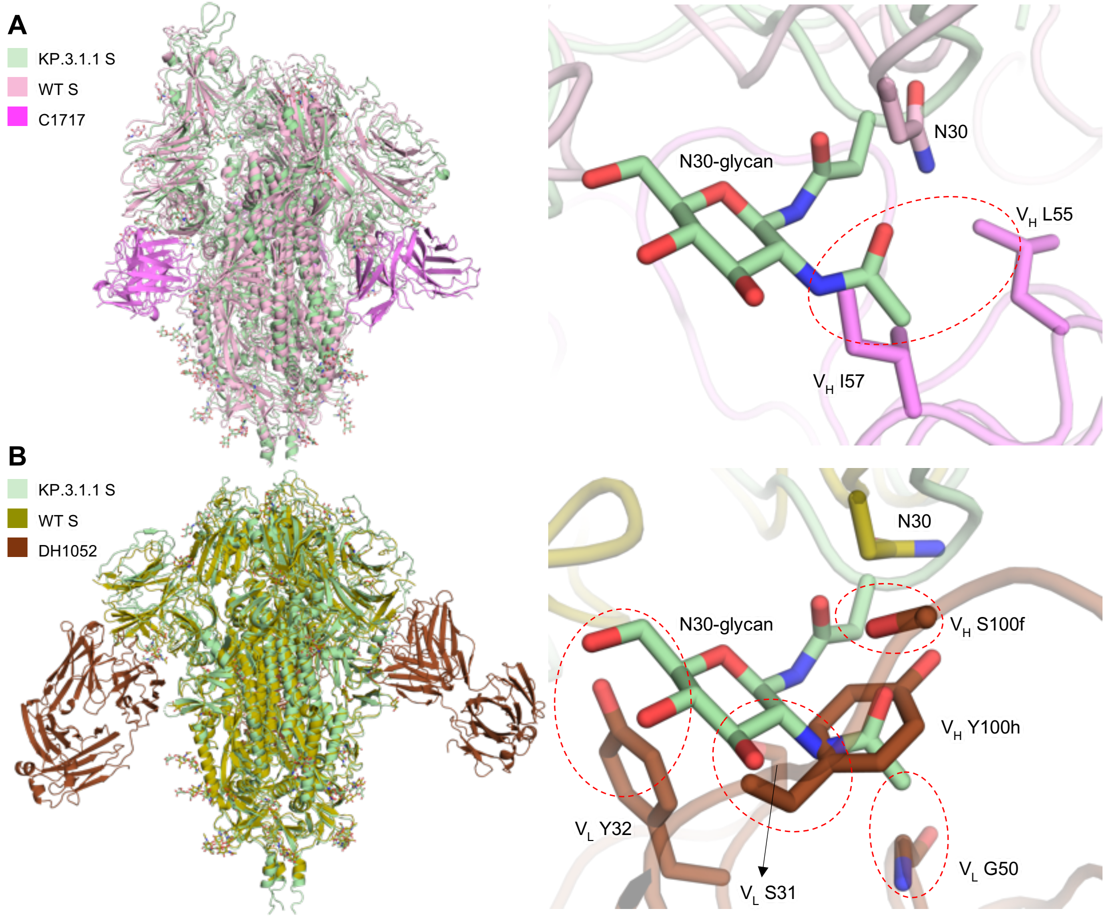


**Figure S9. Structural alignment between KP.3.1.1 S and NTD site vi antibodies. (A)** Left, superimposed structures of KP.3.1.1 S and WT S/C1717 complex. KP.3.1.1 S, WT S, and C1717 are colored in light green, pink, and magenta, respectively. Right, a zoomed-in view of the NTD shows that N30 glycan may clash with C1717 $V_H$ L55 and $V_H$ I57. The red dashed circle highlights the potential clash. **(B)** Right, superimposed structures of KP.3.1.1 S and WT S/DH1052 complex. KP.3.1.1 S, WT S, and DH1052 are colored in light green, olive, and brown, respectively. Right, a zoomed-in view of the NTD shows that N30 glycan may clash with DH1052 $V_H$ S100f, Y100h and $V_L$ S31, Y32, G50. Red dashed circles highlight the potential clashes.

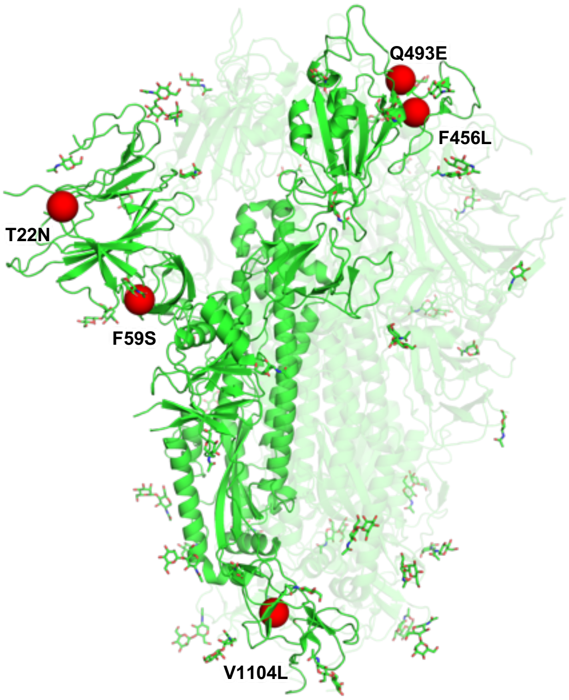


**Figure S10. Mutations (red) on XEC in the context of JN.1 S structure (green) (PDB ID: 8X4H).**

### Table S1. Cryo-EM data collection, processing, and model building statistics.

| Sample | KP.3.1.1 Spike HexaPro + hACE2 | | KP.3.1.1 Spike HexaPro | | | JN.1.11 S31del Q493E Spike HexaPro | | JN.1.11 Spike HexaPro | | JN.1.11 + S31del Spike HexaPro | | JN.1.11.1+S31del Spike HexaPro | |
|---|---|---|---|---|---|---|---|---|---|---|---|---|---|
| **Map** | Local refine RBD+hACE2 | Global refine S with two hACE2 | Local refine RBD+NTD | One RBD Up | All RBD Down | One RBD Up | All RBD Down | One RBD Up | All RBD Down | One RBD Up | All RBD Down | One RBD Up | All RBD Down |
| **EMDB** | 48146 | 48147 | 48148 | 48149 | 48150 | 48151 | 48152 | 48153 | 48155 | 48156 | 48157 | 48158 | 48159 |
| **PDB** | 9ELE | 9ELF | 9ELG | 9ELH | 9ELI | 9ELJ | 9ELK | 9ELL | 9ELM | 9ELN | 9ELO | 9ELP | 9ELQ |
| **Data collection & processing** | | | | | | | | | | | | | |
| **Microscope/ Detector** | Glacios/Falcon4 | | | | | | | | | | | | |
| **Voltage (kV)** | 200 | | | | | | | | | | | | |
| **Magnification** | 190,000 | | | | | | | | | | | | |
| **Recording mode** | Counting | | | | | | | | | | | | |
| **Pixel size (Å)** | 0.718 | | | | | | | | | | | | |
| **Total dose ($e^-/Å^2$)** | 44.84 | | | | | | | | | | | | |
| **Defocus range (μm)** | -0.8 to -1.7 | | | | | | | | | | | | |
| **No. of movie micrographs** | 4334 | | 4336 | | | 4716 | | 3680 | | 4010 | | 7672 | |
| **No. of molecular projection images in map** | 440,016 | 131,224 | 98,127 | 55,445 | 40,554 | 94,590 | 41,085 | 39,569 | 42460 | 63,034 | 59,178 | 53,372 | 112,912 |
| **Symmetry** | C1 | C1 | C1 | C1 | C3 | C1 | C3 | C1 | C3 | C1 | C3 | C1 | C3 |
| **Map pixel size (Å)** | 0.718 | | | | | | | | | | | | |
| **Map resolution (FSC 0.143; Å)** | 3.16 | 2.88 | 3.71 | 2.98 | 2.82 | 2.92 | 2.82 | 2.93 | 2.68 | 2.93 | 2.74 | 2.94 | 2.64 |
| **Map sharpening B-factor ($Å^2$)** | -107.1 | -66.1 | -110.6 | -58.9 | -72.6 | -65.5 | -68.0 | -48.7 | -62.7 | -58.2 | -72.1 | -53.3 | -76.2 |
| **Structure building and validation** | | | | | | | | | | | | | |
| **Model composition** | | | | | | | | | | | | | |
| **Non-hydrogen atoms** | 6529 | 35225 | 4255 | 25512 | 25509 | 25558 | 25299 | 25567 | 25275 | 25446 | 25446 | 23543 | 25479 |
| **Protein residues** | 780 | 4334 | 510 | 3152 | 3153 | 3159 | 3123 | 3157 | 3117 | 3150 | 3150 | 2916 | 3150 |
| **Ligands** | NAG:12/BMA:1 | NAG:58/ BMA:6 | NAG:11 | NAG:52/ BMA:7 | NAG:51/ BMA:6 | NAG:50/ BMA:7 | NAG:51/ BMA:6 | NAG:52/ BMA:7 | NAG:51/ BMA:6 | NAG:49/ BMA:7 | NAG:48/ BMA:6 | NAG:47/ BMA:7 | NAG:51/ BMA:6 |
| **RMSD bond length (Å)/angles (°)** | 0.007/ 1.20 | 0.007/ 1.23 | 0.008/ 1.28 | 0.007/ 1.16 | 0.017/ 1.19 | 0.006/ 1.09 | 0.007/ 1.16 | 0.007/ 1.13 | 0.009/ 1.41 | 0.012/ 1.37 | 0.008/ 1.29 | 0.008/ 1.21 | 0.008/ 1.33 |
| **MolProbity score** | 1.07 | 1.38 | 1.32 | 1.53 | 1.20 | 1.39 | 1.34 | 1.55 | 1.21 | 1.63 | 1.47 | 1.26 | 1.56 |
| **Clash score** | 2.83 | 6.92 | 2.76 | 5.32 | 2.18 | 3.53 | 4.15 | 5.83 | 4.15 | 6.02 | 5.75 | 2.69 | 7.59 |
| **Ramachandran outliers/allowed/favored (%)** | 0/ 1.03/ 98.97 | 0/ 1.94/ 98.06 | 0/ 3.82/ 96.18 | 0/ 3.64/ 96.36 | 0/ 3.32/ 96.68 | 0/ 3.60/ 96.37 | 0/ 2.73/ 97.27 | 0/ 3.57/ 96.43 | 0/ 2.05/ 97.95 | 0.03/ 4.23/ 95.74 | 0/ 2.90/ 97.10 | 0.03/ 3.26/ 96.71 | 0/ 2.80/ 97.20 |
| **Rotamer outliers (%)** | 0 | 0.13 | 0 | 0.04 | 0 | 0 | 0 | 0.14 | 0 | 0.25 | 0.11 | 0.16 | 0.11 |
| **Cβ outliers (%)** | 0 | 0.32 | 0.63 | 0 | 0.1 | 0 | 0 | 0 | 0 | 0.14 | 0.1 | 0.07 | 0.3 |
| **d FSC model (0.5; Å)** | 3.3 | 3.1 | 3.9 | 3.2 | 3.0 | 3.1 | 3.0 | 3.1 | 2.9 | 3.1 | 2.9 | 3.1 | 2.8 |
